# ACCREDIT: A Quality-Aware Agentic Engine for Cell-resolved Cross-modal Image Registration with Dynamic Iterative Tuning

**DOI:** 10.64898/2026.08.08.743602

**Authors:** Le Zhou, Faming Zhao, Tao Ren, Shaun M. Goodyear, Chengyun Tang, Baiyi Li, Tingting Zhang, Yabing Chen, Rosalie C. Sears, Gordon B. Mills, Adel Kardosh, Zheng Xia

## Abstract

Spatial omics across complementary modalities is transforming our understanding of tissue architecture. Realizing this potential requires accurate and robust registration of cross-platform molecular images with hematoxylin-and-eosin (H&E) sections, the primary morphological reference for pathology. Existing methods, however, often fail silently when image orientation is unknown, image contrast is inverted, or tissue overlap is incomplete, producing erroneous registrations without alerting users or attempting recovery. Here, we present ACCREDIT, a quality-aware agentic framework that redefines cross-modal registration as an adaptive decision-making process rather than a one-shot computation. ACCREDIT combines deterministic registration pipelines with a reference-free composite quality score that automatically evaluates registration quality and rejects plausible but biologically incorrect registrations. When registration quality is insufficient, a large language model (LLM)-based rescue agent autonomously diagnoses failure modes and selects targeted recovery strategies, while an optional strategy-learning module captures expert-validated corrections for future reuse. Across Xenium, CODEX, cell-boundary, and IHC-to-H&E registration tasks, ACCREDIT outperformed competing methods by detecting registration failures and improving alignment quality through automated recovery and rescue. Ultimately, ACCREDIT enables robust integration of histology and spatial molecular profiling, providing a foundation for translating spatial omics into routine H&E-based pathology workflows.

## Introduction

Single-cell-resolved spatial-omics and multiplexed imaging platforms, including 10x Xenium, NanoString CosMx and Akoya CODEX^1–5^, now enable in situ measurement of hundreds to thousands of molecular features while preserving tissue architecture^6–8^. Integrating these multimodal measurements has the potential to transform our understanding of tissue architecture with unprecedented resolution and depth. Realizing the full clinical potential of spatial profiling requires integrating molecular measurements with tissue morphology, for which hematoxylin-and-eosin (H&E) staining remains the pathological reference standard. Cross-modal registration to H&E bridges molecular spatial states with the histological context used in routine diagnosis. Consequently, accurate registration of molecular images such as DAPI nuclear morphology, multi-channel protein fluorescence, cell-boundary coordinates and spatial-transcriptomics profiles to corresponding H&E sections is essential for translational multimodal integration. The accuracy and robustness of cross-modal registration directly determine the reliability of downstream spatial analyses. However, current multimodal image-registration methods typically operate as one-shot pipelines that return a single alignment without assessing its quality or providing a self-correcting mechanism to diagnose and recover from registration failures. This limitation is particularly consequential in spatial-omics workflows, where undetected registration errors can compromise downstream biological interpretation^9,10^.

Cross-modal registration in spatial omics is challenging for three major reasons. First, image orientation is often unknown, as sample loading and scanning can introduce arbitrary flips and 90-degree rotations, causing methods that assume approximate pre-alignment to fail silently^11–14^. Second, image appearance differs fundamentally across modalities: fluorescence images typically present bright signal on a dark background, whereas H&E and immunohistochemistry (IHC) images show dark tissue structures on a bright field, which weakens the intensity-similarity assumptions underlying many registration metrics. Third, the fields of view are often only partially overlapping, because CODEX, Visium and some Xenium acquisitions may capture only a subset of the matched H&E section. Under these conditions, global optimization can converge to degenerate solutions that maximize apparent overlap while sacrificing anatomical correspondence. Together, these challenges suggest that robust cross-modal registration requires more than an alignment pipeline alone; it also requires a framework that can disambiguate orientation, accommodate cross-modal appearance mismatch, and detect implausible solutions^15–17^.

Although many registration methods address one or more of these issues, most were designed under narrow assumptions and do not provide a unified mechanism for quality assessment, failure diagnosis, and adaptive recovery. Group-wise serial-section approaches such as VALIS^18^ perform well on same-stain benchmarks^19^ but typically assume approximate pre-alignment and do not recover when image orientation is incorrect. Deep-learning deformable approaches, including VoxelMorph^20^, TransMorph^21^ and DHR^22^, the ACROBAT 2023 winner^23^, can achieve strong local refinement once a reasonable initialization is available, but they do not explicitly search over flips or large rotations. Same-section mosaicking and multiplexed-imaging frameworks such as Palom^24^ and MCMICRO^25^ generally assume a shared tissue section and pixel scale, whereas platform-specific tools such as VoltRon^13^ and Xenium-Align^26^ provide narrow modality coverage. Although classical registration toolkits including ANTs^27^, elastix^28^ and SimpleITK^29^ offer powerful numerical components, they can still fail in challenging registration tasks. In summary, existing registration methods address individual aspects of these challenges but do not jointly resolve them, particularly when multiple failure modes coexist within a single registration task. When registration fails, the underlying cause can vary, including incorrect orientation, modality-descriptor mismatch, partial overlap and local deformation, and each failure mode requires a different recovery strategy. Consequently, no fixed compute-once pipeline can reliably diagnose and recover from all failure modes. Robust cross-modal registration therefore requires runtime quality assessment coupled with adaptive recovery during the registration process.

Here we present ACCREDIT (Agentic Cell-resolved Cross-modal Registration Engine with Dynamic Iterative Tuning), a quality-aware agentic registration framework that embeds LLM-based reasoning as a runtime decision component within a closed-loop scientific image-registration system. Unlike existing LLM-assisted biology tools, in which the agent typically operates outside the algorithm to help users write code, select parameters or plan workflows, ACCREDIT places the agent inside the registration loop. Deterministic registration modules perform the numerical alignment, a reference-free Composite Quality Score (QCS) evaluates each candidate output and decomposes failure signals, and the LLM agent uses these diagnostics to infer likely failure modes and select the next recovery strategy. A complementary, optional human-guided experience library stores user-validated recovery strategies so that successful corrections can be automatically reused in future runs. This design establishes a generalizable pattern for scientific computing problems with large combinatorial strategy spaces: deterministic algorithms perform the numerical heavy lifting, agentic reasoning performs failure diagnosis and strategy selection, and an optional experience-learning layer accumulates user-validated strategies across cases. To our knowledge, ACCREDIT is the first image-registration framework to explicitly embed LLM-based agentic reasoning as a runtime decision component within a quality-gated registration loop. We evaluated ACCREDIT on DAPI-to-H&E registration across public Xenium samples, cell-boundary-to-H&E registration as an automated alternative to the Xenium Explorer workflow, and IHC-to-H&E registration on pancreatic cancer samples, and further examined whether improved registration facilitates downstream single-cell and cross-modal spatial analyses.

## Results

### 1. Overview of the ACCREDIT framework

The ACCREDIT framework operates as a quality-gated, agentic registration cascade with automated modality routing, QCS-based acceptance and an adaptive rescue and strategy-learning module (**Fig. 1**). Inputs are first classified by modality, routed to a modality-specific deterministic registration pipeline and evaluated by a reference-free composite quality score (QCS; Methods, “Composite Quality Score”). Outputs exceeding the QCS acceptance threshold are returned directly, whereas lower-scoring outputs are converted into structured diagnostic feedback through the QCS decomposition. In the autonomous rescue loop, the LLM agent does not directly estimate image transformations. Instead, it uses the QCS components, coverage penalties, modality metadata and previous pipeline configuration to diagnose likely failure modes and select recovery strategies such as wider orientation search, enhanced preprocessing, channel re-selection, ROI-scale search or alternative registration backends. Each selected strategy is executed by the deterministic backend and re-evaluated by QCS.

**Fig. 1.**
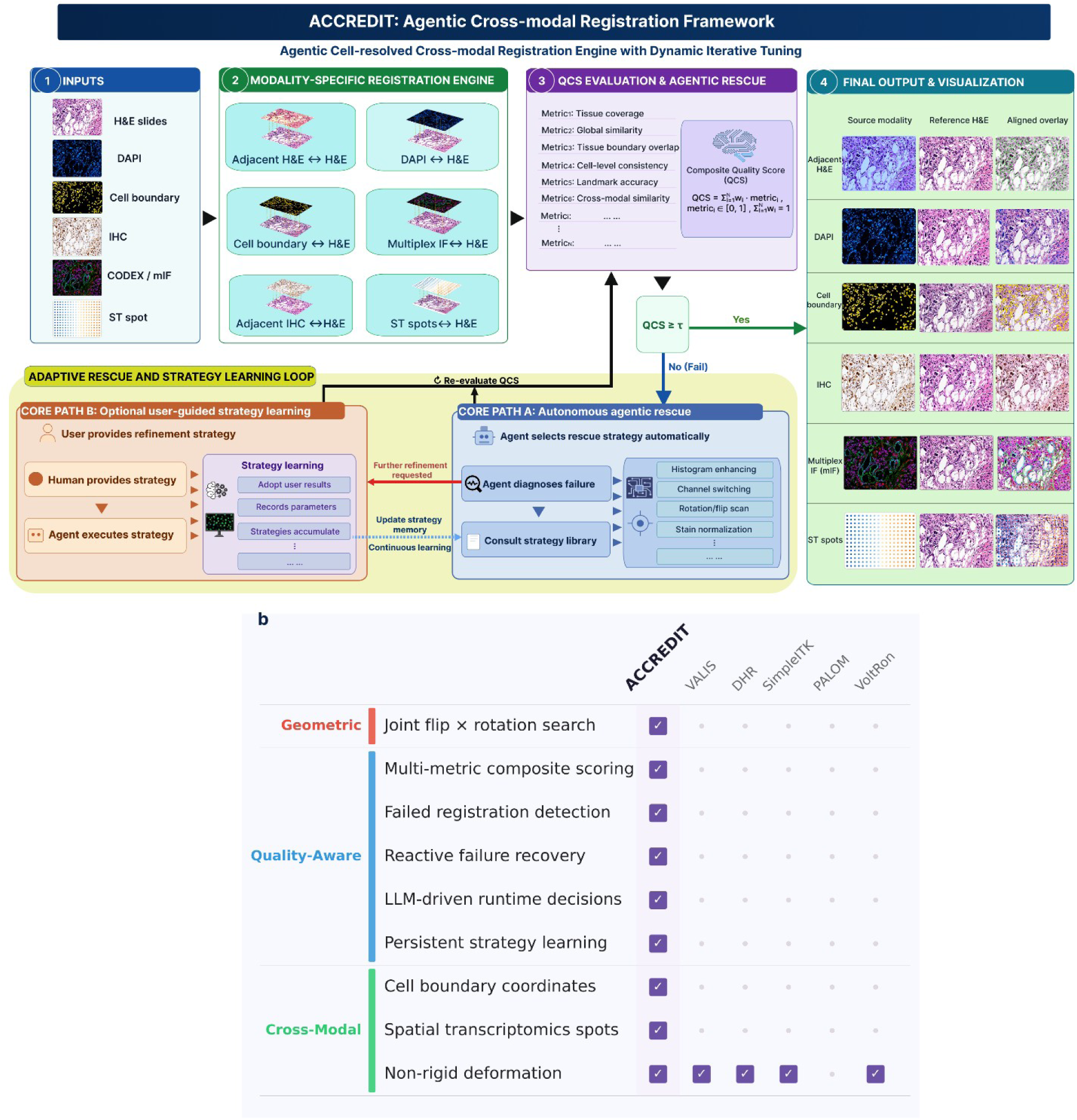
ACCREDIT framework and capability comparison. a, Entry-point modality detection classifying each input into one of six modality pairings. **b,** Modality-specific routing to the corresponding deterministic registration pipeline. **c,** Composite quality score (QCS) evaluation of each registration output. **d,** Return of the best-scoring acceptable registration result. The adaptive rescue and strategy-learning loop contains two paths. **Core Path A,** autonomous agentic rescue triggered when the default registration output falls below the task-specific QCS threshold; the agent diagnoses failure modes and selects recovery strategies from a predefined strategy library while deterministic modules perform the image transformation. **Core Path B,** optional user-guided strategy-learning path, in which user-provided or user-validated refinement strategies are abstracted into reusable rules and stored in the experience knowledge base. A detailed DAPI-to-H&E example is shown in Supplementary Fig. 1. **e,** Capability matrix comparing ACCREDIT against five existing tools (VALIS^18^, DHR, SimpleITK, PALOM and VoltRon^13^); filled cells indicate that the method provides the listed capability.

For routine use, ACCREDIT is designed to operate automatically: inputs are routed by modality, evaluated by QCS and, when necessary, improved through autonomous agentic rescue without requiring user intervention. In addition, ACCREDIT provides an optional user-guided strategy-learning path for cases in which a user wishes to further improve an automated result or convert a validated correction into reusable pipeline knowledge. In this path, the user can inspect an intermediate output, request further refinement or provide a refinement strategy. The agent then abstracts the validated correction into a reusable rule stored in the experience knowledge base. **Supplementary Fig. 1** illustrates this design using separate formalin-fixed paraffin-embedded (FFPE) sections of breast cancer stained with DAPI and H&E, respectively (Breast Cancer FFPE Rep1). After automated DAPI-to-H&E registration and autonomous rescue, residual local mismatch remained after automated improvement. User-guided refinement reused the intermediate warped DAPI image and resized H&E reference for second-pass matching and residual-offset correction. The resulting correction was abstracted into a reusable rule for re-matching intermediate outputs when residual local mismatch persists after rescue and was incorporated into the updated DAPI-to-H&E pipeline. The learned rule could then be applied automatically to subsequent samples, such as Breast Cancer FFPE Rep2, without repeated human intervention. Thus, the agentic contribution of ACCREDIT is not external workflow assistance, but runtime failure diagnosis, autonomous strategy selection and optional conversion of user feedback into reusable registration knowledge.

### 2. ACCREDIT improves DAPI-to-H&E benchmark across Xenium samples

Applying ACCREDIT’s DAPI-to-H&E pipeline (Methods, “DAPI-to-H&E pipeline”) to 16 public Xenium samples spanning 13 tissue types and both Xenium V1 and Xenium Prime chemistries yielded a mean composite quality score (QCS) of 0.77 with zero silent failures, making ACCREDIT the only method to successfully process every sample. Fig. 2a shows the per-sample bubble grid, with mean QCS values in descending order: ACCREDIT (0.77), VALIS (0.67), DHR (0.59), SimpleITK (0.35), VoltRon (0.25), and Palom (0.16). The grid also shows that ACCREDIT’s worst sample (QCS = 0.65 on Prostate) still exceeds the mean QCS of four competing methods. Crosses (x) mark silent failures, which cluster in Palom and VoltRon on partial overlap or cross-platform cases.

**Fig. 2.**
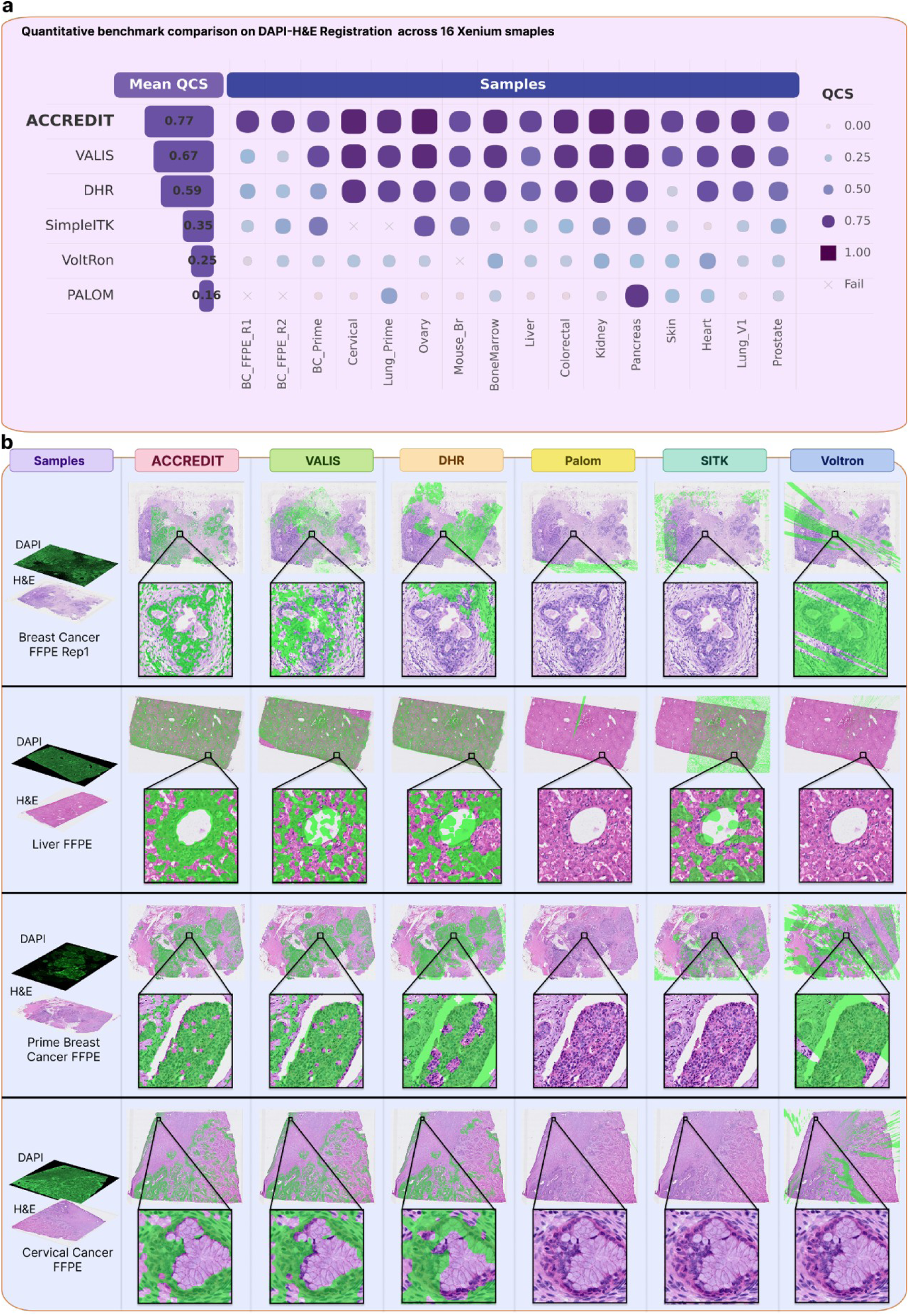
DAPI-to-H&E benchmark across 16 public Xenium samples. **a**, Per-sample composite quality score (QCS) bubble grid for ACCREDIT and five competing methods (VALIS, DHR, SimpleITK, VoltRon, Palom) evaluated on 16 Xenium samples spanning 13 tissue types and both Xenium V1 and Xenium Prime chemistries. The first column shows mean QCS per method; subsequent columns show per-sample QCS with bubble size proportional to QCS; crosses (×) mark silent failures. **b**, Qualitative DAPI-to-H&E overlay comparison on four representative samples (Breast Cancer FFPE Rep1, nondiseased Liver FFPE, Prime Breast Cancer FFPE, Cervical Cancer FFPE) spanning the difficulty spectrum. Each row shows the input DAPI and matched H&E (left) followed by overlaid registration outputs of the six methods (right), with triangular insets zooming into a tissue region. Overlays render DAPI in green and H&E in magenta; co-located signal appears purple, while misregistration leaves green and magenta visibly separated.

A qualitative comparison of four representative samples spanning the difficulty spectrum (**Fig. 2b**) shows that ACCREDIT consistently places DAPI signals on the corresponding H&E structure where competing methods do not. Breast Cancer FFPE Rep1 is the most demanding case because it combines an instrument-induced flip with partial field-of-view overlap. ACCREDIT recovers both the orientation and the correct tissue placement, whereas competing methods either retain the mirrored orientation or collapse into a degenerate corner solution. On nondiseased liver FFPE and Prime Breast Cancer FFPE, the main difference is tissue-boundary precision, with ACCREDIT placing DAPI and H&E contours into closer register than the other methods. On Cervical Cancer FFPE, the zoom inset provides cell-level confirmation, with ACCREDIT aligning DAPI and H&E nuclei more closely than any competing overlay.

Per-method failure patterns point to distinct but interpretable limitations. VALIS and DHR are strongest when a roughly correct initialization is already available, but both are vulnerable when the acquisition introduces a flip or large rotation. Breast Cancer FFPE Rep1 is the clearest example. Against the SimpleITK baseline (0.77 vs 0.35), which represents the strongest pipeline assembled from the bare numerical engine alone, ACCREDIT layers 96-candidate search, QCS-driven decisions and rescue tools above the same engine. The 0.42-point gap therefore quantifies the contribution of the upper-layer architecture rather than any difference in the underlying numerical library. VoltRon and Palom struggle most on cross-modal descriptor mismatch or cross-platform scale mismatch. These patterns indicate that the performance gap reflects design envelope rather than a single isolated algorithmic component.

Taken together, the per-sample bubble grid (**Fig. 2a**), the representative overlays (**Fig. 2b**) and the per-competitor summary show that ACCREDIT’s advantage lies in handling the combined challenge of unknown orientation, cross-modal appearance mismatch and partial overlap within a single framework. Breast Cancer FFPE Rep1 makes this concrete in a single image: it is the only sample in the 16-sample benchmark on which the instrument introduced both a flip and a partial-field-of-view condition, and it is the sample on which every competing method fails in a visibly distinct way: VALIS and DHR leave the overlay mirrored because their initialization grids do not consider flip, Palom and VoltRon collapse the warped tissue into a degenerate corner that maximizes pixel overlap at the cost of anatomical correspondence, and SimpleITK retains gross orientation but offsets the tissue out of register. ACCREDIT is the only method that correctly recovers both the orientation and the partial-overlap registration, and does so as part of the routine output of the pipeline rather than after manual intervention.

Beyond the Xenium benchmark shown in **Fig. 2**, we further evaluated ACCREDIT on a glioma CODEX-to-H&E dataset to assess cross-platform generalization. In the combined 28-sample benchmark, including 16 Xenium DAPI-to-H&E pairs and 12 glioma CODEX-to-H&E pairs, ACCREDIT showed the highest mean QCS and a compact high-performance distribution across both datasets, whereas competing methods showed broader variability and more dataset-dependent performance (**Supplementary Fig. 2**).

### 3. ACCREDIT enables automated cell-boundary-to-H&E registration

ACCREDIT’s cell-boundary pipeline (Methods, “Cell-boundary-to-H&E pipeline”) automatically aligns Xenium per-cell boundary coordinates to the matched H&E section in under five minutes per sample (mean 4.8 min per sample, range 1–10 min), through a four-stage workflow with an automatic rescue path when the initial feature-based alignment is insufficient (**Fig. 3a**).

**Fig. 3.**
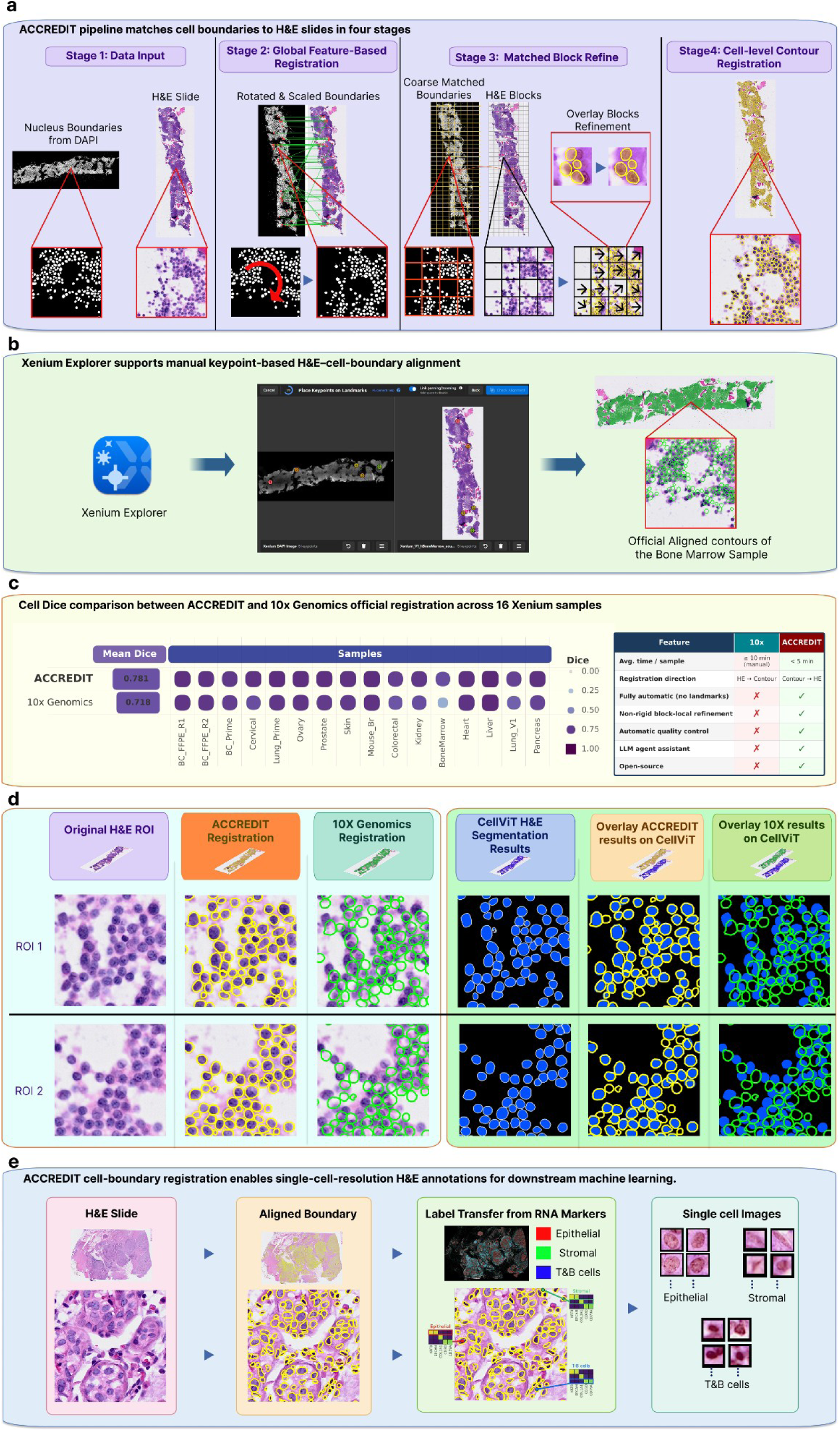
Cell-boundary pipeline replacing the 10x Xenium Explorer workflow, third-party validation and downstream single-cell application. **a**, ACCREDIT’s four-stage cell-boundary pipeline (Methods, “Cell-boundary-to-H&E pipeline”). The panel includes both the stage schematic and a summary table indicating the role of each step, from coordinate conversion and global affine initialization to local refinement, final polygon transformation and the optional rescue path. **b**, The 10x Genomics Xenium Explorer GUI workflow that ACCREDIT replaces. **c**, Per-sample cell Dice comparison between ACCREDIT and the 10x official alignment on the 16 Xenium samples from Fig. 2, together with a workflow-property table contrasting the two approaches in terms of automation, batch processing, quality control and alignment direction, highlighting that ACCREDIT maps contours into H&E image space whereas 10x projects H&E into contour space. **d**, Two ROIs (BoneMarrow and Ovarian) showing per-cell comparison with CellViT as a third-party H&E-only cell-segmentation reference. Each row shows the original H&E ROI, ACCREDIT-aligned boundary, 10x-aligned boundary, CellViT segmentation, and ACCREDIT/CellViT and 10x/CellViT overlays. e, Downstream label-transfer application: RNA-derived cell-type labels transferred from the spatial-omics layer onto matched H&E coordinates, producing single-cell H&E image patches sorted by type.

ACCREDIT replaces the 10x Genomics’ Xenium Explorer graphical user interface (GUI) workflow (**Fig. 3b**). Xenium Explorer requires either manual selection of corresponding keypoints in paired H&E and DAPI images or loading of a pre-computed alignment CSV provided only for demonstration datasets. The manual route is slow, requires user expertise, and provides no internal quality assessment, whereas the official-CSV route is unavailable for user-generated samples and does not support batch processing. Essentially, the 10x workflow projects the H&E image into boundary-coordinate space, effectively matching a high-dimensional histology image to a lower-dimensional contour representation. In contrast, ACCREDIT takes the opposite approach by aligning boundary coordinates into the H&E image space, which is more natural for morphology-based validation and downstream image analysis. The workflow-comparison summary in **Fig. 3c** makes these differences explicit by contrasting ACCREDIT and 10x in terms of automation, batchability, quality control and alignment direction. ACCREDIT replaces both routes with a single QCS-gated batch workflow that runs without user intervention.

On the same 16 Xenium samples used in Fig. 2, ACCREDIT outperformed the official 10x alignment on cell Dice for every sample (Fig. 3c; mean cell Dice 0.78 vs 0.72; per-sample win rate 16/16), registering a total of 4,020,268 cells. The largest gain was observed on BoneMarrow (0.68 vs 0.38; Dice improvement = 0.30), an especially dense and morphologically complex sample on which the 10x alignment visibly degraded, while ACCREDIT’s local refinement remained stable; the smallest gains, such as those for Liver (Dice improvement = 0.0030) and Heart (Dice improvement = 0.0040), occurred on samples where the baseline alignment was already relatively strong. Fig. 3c also includes a workflow-property comparison showing that ACCREDIT provides automatic batch processing and internal quality control, whereas 10x Xenium Explorer depends on manual interaction or an external pre-computed alignment file and registers in the opposite direction by projecting H&E into contour space rather than mapping contours into image space.

Because cell Dice is not itself a ground-truth biological label, we used CellViT^30^, an H&E-only cell-segmentation model that predicts per-cell instance boundaries with no knowledge of either the Xenium boundary file or the 10x alignment CSV, as a third-party reference (**Fig. 3d**). ACCREDIT’s boundaries match CellViT’s H&E-derived segmentation at sub-cell resolution, supporting the interpretation that the improved Dice is not just a metric artifact but reflects genuinely enhanced cell-level spatial registration.

This level of boundary alignment enables a downstream single-cell H&E analysis pipeline (**Fig. 3e**). After registration, RNA-derived cell-type labels can be transferred from the spatial-omics layer onto H&E coordinates to generate single-cell image patches sorted by type. Because even a one-to two-cell-diameter offset can misassign labels, this application depends directly on accurate and failure-aware boundary registration.

Together, the 16/16 cell-Dice win (**Fig. 3c**), the CellViT-based external validation (**Fig. 3d**) and the downstream label-transfer example (**Fig. 3e**) show that ACCREDIT replaces the manual Xenium Explorer workflow with an automated and more accurate alternative. The single-cell H&E labeling workflow enabled by ACCREDIT cell-boundary registration is detailed in **Supplementary Fig. 3**.

### 4. ACCREDIT generalizes to IHC-to-H&E registration

We evaluated the ACCREDIT-IHC hybrid backend on representative pancreatic adenosquamous carcinoma samples using the fixed recipe described in Methods, “IHC-to-H&E hybrid backend”, without per-sample parameter selection. **Fig. 4a** shows qualitative overlays for two samples, referred to as Sample 1 and Sample 2. In both examples, ACCREDIT placed the IHC signal in closer register with the matched H&E tissue structure than the competing methods, particularly in the zoomed regions where local tissue boundaries and high-intensity IHC regions are more clearly aligned. **Fig. 4b** provides a detailed metric-level comparison for the two displayed samples. On Sample 1, ACCREDIT achieved the highest QCS of 0.92, compared with 0.66 for DHR, 0.45 for VALIS, 0.10 for VoltRon, 0.03 for SimpleITK and 0.02 for Palom. On Sample 2, ACCREDIT again achieved the highest QCS of 0.86, followed by VALIS at 0.77, DHR at 0.65, VoltRon at 0.21, SimpleITK at 0.05 and Palom at 0.00. Across both representative cases, ACCREDIT showed stronger tissue overlap, boundary agreement and coverage-related performance, whereas competing methods showed residual local mismatch, reduced feature consistency or unstable deformation in the qualitative overlays.

**Fig. 4.**
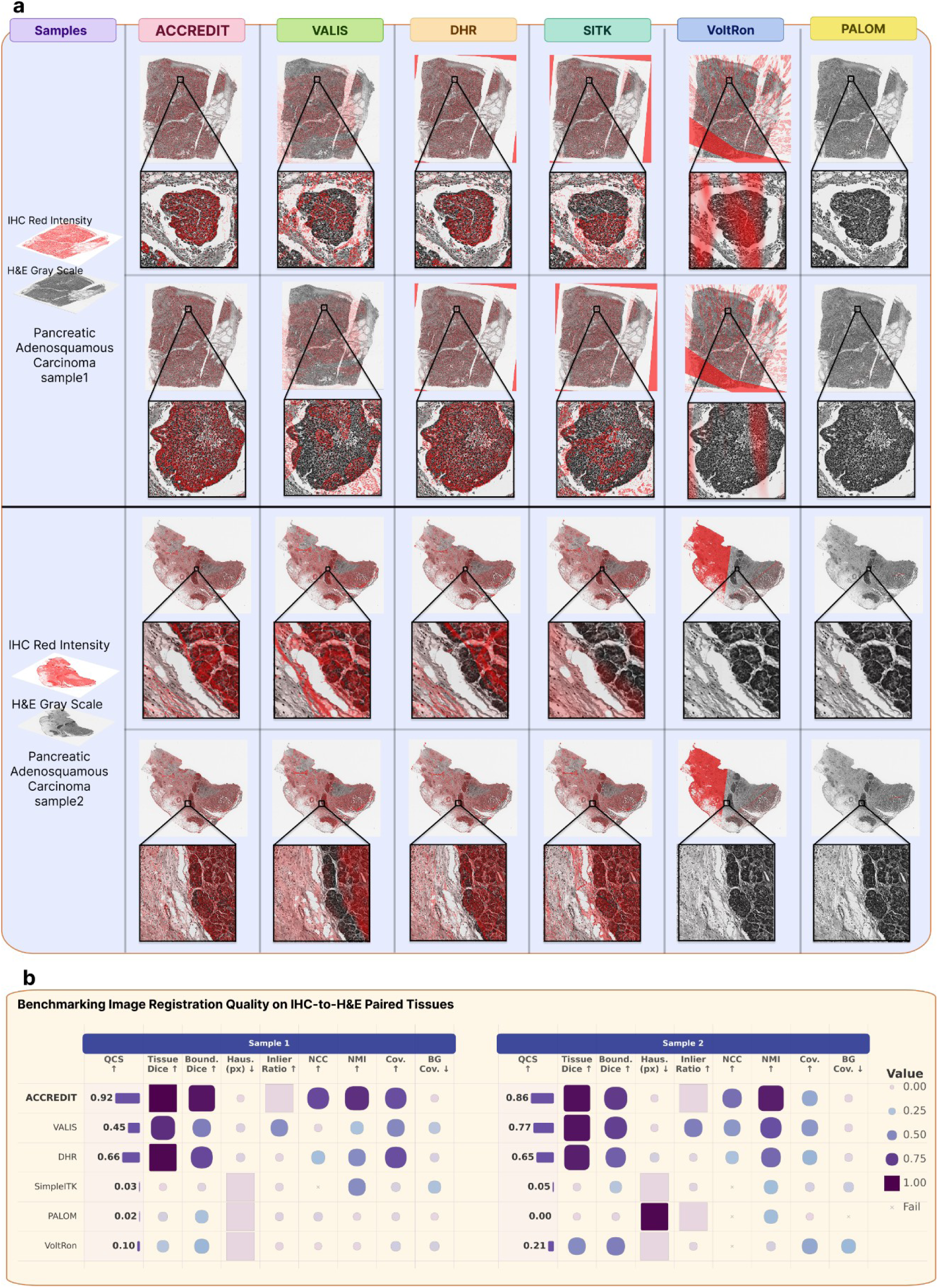
IHC-to-H&E registration benchmark on representative pancreatic adenosquamous carcinoma samples. **a**, Qualitative IHC-to-H&E overlay comparison on two representative pancreatic adenosquamous carcinoma samples, referred to as Sample 1 and Sample 2. For each sample, the input IHC intensity image and matched H&E grayscale image are shown on the left, followed by overlaid registration outputs from ACCREDIT and five competing methods: VALIS, DHR, SimpleITK, VoltRon and Palom. The IHC signal is rendered in red and overlaid on the H&E grayscale background to visualize spatial correspondence. Zoomed regions highlight local registration differences across methods. **b**, Quantitative metric comparison for the two representative samples shown in **a**. For each method, the bubble grid reports QCS together with individual evaluation metrics, including tissue Dice, boundary Dice, Hausdorff distance, inlier ratio, normalized cross-correlation, normalized mutual information, tissue coverage and background coverage. Bubble size and color intensity indicate metric value; crosses indicate failed or unavailable metric values.

### 5. Multimodal spatial alignment enables cross-modal characterization of cellular neighborhoods

Spatial protein imaging (e.g., CODEX) enables robust identification of cellular neighborhoods (CNs) that capture higher-order tissue organization, but these structures are often difficult to interpret at the molecular level due to the lack of matched transcriptomic information^31,32^. Conversely, spatial transcriptomics (e.g., Xenium) provides rich gene expression data but lacks reliable protein-based definitions of spatial neighborhoods^10^. To bridge this gap, we performed physical alignment of adjacent Xenium and CODEX tissue sections, enabling direct mapping of protein-defined CNs onto transcriptomic space (**Fig. 5a**). The multimodal datasets were obtained from the SPATCH database^10^, and cell-type annotations were adopted from the original study.

**Fig. 5.**
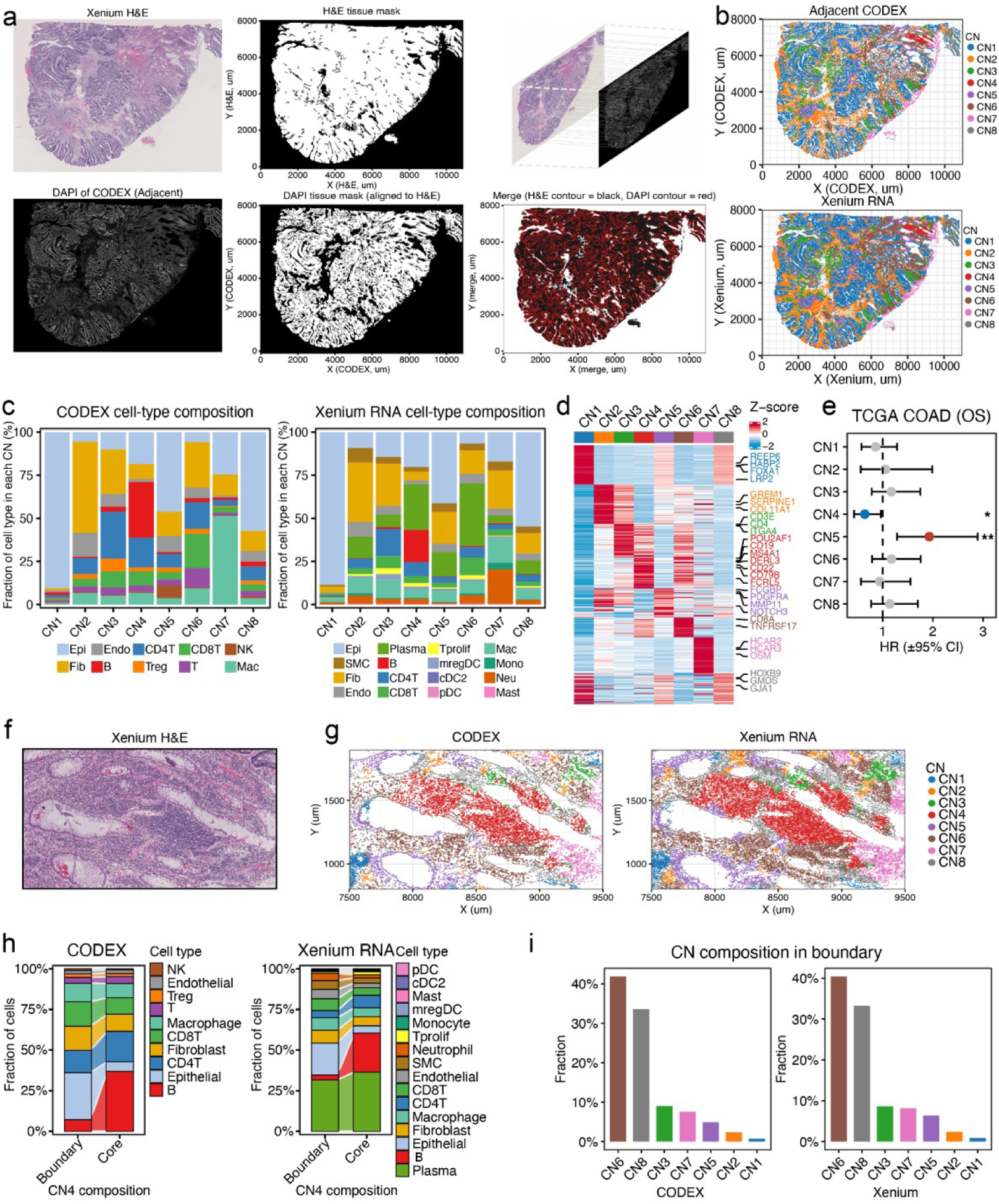
Multimodal cellular neighborhood characterization across CODEX and Xenium. **a**, Physical alignment of adjacent Xenium and CODEX tissue sections, shown with matched H&E and CODEX DAPI images, derived tissue masks and their overlay. **b**, Spatial distribution of cellular neighborhoods (CNs) identified from CODEX protein data and projected onto Xenium coordinates after alignment. **c**, Cell-type composition of each CN derived from CODEX protein data and Xenium RNA data. **d**, Differential expression signatures across CNs based on Xenium RNA data. **e**, Association of CN-linked states with overall survival in the TCGA COAD cohort using MuSiC deconvolution and Cox proportional hazards models. **f**, Representative H&E image illustrating a secondary follicle-like region. **g**, Zoomed views showing concordant CN architecture across CODEX and Xenium. **h**, Cell-type composition of CN4 in boundary versus core regions. **i**, Distribution of additional CN states within the boundary region of the SFL-like niche across both modalities.

Alignment based on H&E and CODEX DAPI images showed high concordance between tissue masks, demonstrating accurate spatial registration (**Fig. 5a**). Using CODEX protein data and corresponding cell-type annotations (see Methods), we identified eight distinct CNs (CN1-CN8) that capture higher-order spatial organization (**Fig. 5b** and **Fig. S4a**). Because CODEX and Xenium cells were co-registered in a shared H&E coordinate system, each Xenium cell was assigned the CN label of its nearest CODEX neighbor, thereby projecting protein-defined CN states into the RNA space (see Methods). Following this mapping, CN distributions in Xenium closely recapitulated those observed in CODEX (**Fig. 5b**). Although the original cell-type annotations differed between the CODEX and Xenium datasets due to modality-specific annotation schemes^10^, the overall cell-type composition and spatial organization within each CN remained highly consistent across modalities (**Fig. 5c**). Both global and local annotation patterns further supported the robustness of cross-modal mapping (**Fig. S4b-c**).

This cross-modal mapping enables direct assignment of transcriptomic programs to protein-defined spatial neighborhoods. Leveraging Xenium RNA data, we systematically characterized these programs across CNs, revealing distinct gene expression signatures for each neighborhood (**Fig. 5d**), enabling molecular annotation of protein-defined spatial states. Notably, CN4 was enriched for B cell-associated features^33^, including canonical markers such as *CD79B*, *MS4A1*, and *CD22*. In contrast, CN5 exhibited enrichment of stromal and extracellular matrix remodeling programs^34^, including genes such as *PDGFRA*, *MMP11*, *NOTCH3*, and *FCGBP*. Spatially, CN5 was preferentially localized at the tumor-stroma interface, consistent with a niche characterized by active-matrix remodeling and invasive potential. To assess clinical relevance, CN-associated proportions were estimated in bulk transcriptomic data using MuSiC deconvolution with Xenium RNA as reference^35^, followed by Cox proportional hazards analysis in the TCGA COAD cohort ^36^. This analysis revealed that specific CN states were associated with patient outcomes, with CN4 linked to improved prognosis and CN5 associated with adverse outcomes (**Fig. 5e**).

Inspection of the aligned H&E image revealed a localized region with lymphoid-like morphology, consistent with a secondary follicle-like (SFL-like) structure^37^ (**Fig. 5f**). Mapping CN labels to this region showed that it was predominantly occupied by CN4, suggesting that this CN represents a spatially coherent immune niche. Zoom-in views across CODEX and Xenium data demonstrated concordant CN architecture within this region (**Fig. 5g**). Further spatial stratification revealed a clear core-boundary organization within this SFL-like niche. The core region was enriched for B cells and plasma cells, whereas the boundary exhibited increased heterogeneity, including T cells, macrophages, and stromal components (**Fig. 5h**). This spatial organization was consistently observed across both protein and RNA modalities, supporting its biological validity. In addition, epithelial enrichment in the boundary suggests that this niche is positioned at the tumor-immune interface rather than within tumor cores. Analysis of CN distribution within the boundary further revealed enrichment of multiple CN states with consistent patterns across modalities (**Fig. 5i**), highlighting the compositional complexity of the niche periphery. Together, these findings demonstrate that multimodal spatial alignment enables identification of biologically meaningful immune niches with structured spatial organization and potential clinical relevance.

## Discussion

Cross-modal image registration in spatial omics is often treated as a compute-once operation: run an algorithm, accept the output, and proceed to downstream analysis. Our results instead support a quality-gated view in which candidate registrations must be evaluated before they are trusted. The clearest practical consequence is reduced silent misregistration under unknown orientation, which was the dominant failure mode across competing methods in the DAPI-to-H&E benchmark assessed herein. When imaging instruments introduce flips or 90° rotations, methods that assume approximate pre-alignment can produce plausible looking but biologically incorrect overlays. By explicitly searching orientation hypotheses and selecting among them with QCS, ACCREDIT removes this assumption and broadens the range of cases that can be reliably handled, particularly in an automated batch approach that improves speed and reliability. At the same time, this design is complementary to, rather than a replacement for specialized deformable methods: when orientation is already known, those methods can still be effective for local refinement and can be incorporated as modular backends.

The composite quality score (QCS) addresses a shared weakness of static registration tools: most do not evaluate their own output. QCS combines five complementary signals, including tissue-mask overlap, boundary-band Dice, normalized Hausdorff-distance score, feature-inlier ratio and normalized mutual information, together with coverage penalties that suppress degenerate solutions. No single metric is sufficient on its own: tissue Dice can remain moderate despite boundary-level misalignment, whereas NMI can remain high in locally homogeneous regions despite gross spatial offset. Their combination reduces these blind spots, as illustrated by the DAPI-to-H&E benchmark used herein, where visually implausible alignments are penalized despite superficially reasonable overlap. More broadly, these results suggest that reference-free quality assessment should be treated as a core component of cross-modal biomedical registration rather than as a post hoc diagnostic.

The IHC-to-H&E benchmark further illustrates the value of ACCREDIT’s modular design. Adjacent serial sections stained with different protocols share gross tissue structure but differ in local contrast and often exhibit non-rigid deformation introduced during sectioning, mounting and staining. On this task, a deterministic hybrid recipe that combines robust global initialization with successive non-rigid refinement achieved strong performance without per-sample tuning and without invoking additional rescue stages in the representative IHC cases. This outcome suggests that the framework can accommodate task-specific deformation patterns by pairing a common quality-control layer with a modality-appropriate registration backend. At the same time, broader validation across additional IHC stains and tissue types will be needed to determine how far this fixed recipe generalizes.

The cross-modal capability of ACCREDIT rests on a biological rather than purely computational insight: the hematoxylin component of H&E and the DAPI fluorescent stain both reflect nuclear density, even though they appear visually dissimilar. Extracting a hematoxylin-emphasizing representation therefore converts a difficult cross-modal matching problem into one involving two views of a shared biological substrate. The same principle extends to multi-channel protein imaging, where QCS-guided channel selection identifies the channel whose tissue pattern best supports alignment for a given sample. The practical value of this strategy is reflected in the downstream analysis in **Fig. 5**, where physical alignment between CODEX and Xenium enabled transfer of protein-defined neighborhood structure into transcriptomic space. In this sense, ACCREDIT is complementary to computational integration frameworks such as MaxFuse, because it provides the spatial correspondence layer on which multimodal interpretation can depend^38^.

Several limitations should be noted. First, although ACCREDIT is designed to support six spatial-omics modalities, the present study systematically benchmarks DAPI-to-H&E and cell-boundary-to-H&E registration, while using representative IHC-to-H&E cases to demonstrate extension to serial-section immunohistochemistry; broader validation on additional platforms requires further evaluation and possibly modality-specific retuning. Second, tissue-processing artifacts between adjacent serial sections, such as tearing, folding and differential shrinkage, are only partially addressed by the current non-rigid backends. Third, the LLM rescue agent depends on external API access and adds computational cost, even though rescue was not required in the representative IHC-to-H&E cases, and most high throughput use cases are expected to remain in the deterministic first stage. Fourth, QCS is still a proxy metric: although the CellViT comparison supports its biological relevance at the cell level, extreme tissue morphology or imaging artifacts may still mislead it. These limitations define the main boundaries of the current framework and the most important directions for external validation.

The cell-boundary results also point to applications beyond registration itself. Once per-cell contours are accurately mapped into H&E image space, they provide a natural scaffold for downstream single-cell image analysis, including morphology-based cell typing, image-derived molecular prediction and neighborhood analysis in frameworks such as Squidpy and Giotto. Future work could further improve cross-modal matching by using stain-invariant representations from emerging pathology foundation models. More generally, the learning mechanism built into ACCREDIT suggests a path toward progressively expanding validated registration strategies across tissue types and imaging settings ^39–42^.

In summary, ACCREDIT frames cross-modal spatial-omics registration as a quality-aware decision process rather than a fixed one-shot pipeline. Across DAPI-to-H&E, cell-boundary-to-H&E and IHC-to-H&E evaluations, the framework reduced silently propagated misregistration by combining explicit quality control, adaptive recovery and modular task-specific backends. More broadly, ACCREDIT illustrates a general agentic design pattern for scientific image analysis: deterministic algorithms perform the numerical heavy lifting, QCS exposes interpretable failure signals, an embedded LLM agent performs runtime failure diagnosis and autonomous strategy selection, and an optional experience library accumulates user-validated recovery paths for future automated reuse. This differs from existing LLM-assisted biology systems in which the agent typically remains outside the algorithm as a workflow planner or coding assistant. By embedding agentic reasoning inside the quality-gated registration loop, ACCREDIT suggests a broader framework for self-correcting scientific image-analysis systems beyond spatial omics registration.

## Methods

### Mathematical notation and system overview

ACCREDIT frames cross-modal registration as a quality-gated adaptive registration system in which deterministic registration modules estimate transforms, QCS evaluates candidate outputs, and the agent performs failure-mode diagnosis and strategy routing when the default pipeline is insufficient. Formally, let the moving image, the H&E reference image, their tissue masks and the reference-image domain be defined as *I_m_, I _r_, M_m_, M_r_*. A registration attempt estimates a transform from moving coordinates into the H&E reference coordinate system and produces a warped moving image:

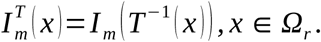

Here, *I_m_*is the moving image before registration, *I _r_* is the H&E reference image, T is the spatial transform from moving coordinates into reference coordinates, and *Ω_r_* is the reference-image domain. The superscript T means that the moving image has been warped by T. In practical terms, this notation defines what the registered moving image looks like after applying the estimated transform. Rather than accepting the output of a single fixed algorithm, ACCREDIT maintains a finite library of predefined recovery strategies, but these strategies are not exhaustively enumerated during routine execution. Instead, candidate transforms generated by the default cascade and any triggered recovery modules are evaluated by QCS, and the final output is selected as the highest scoring acceptable result among attempted configurations.

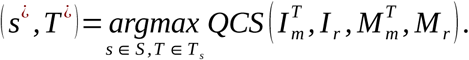

For notational simplicity, *QCS* (*T*) denotes the composite quality score computed from the registration output produced by transform T, including the warped moving image, warped moving mask and corresponding H&E reference measurements.

The implemented workflow separates numerical optimization from strategy-level decision making. First, modality-specific deterministic registration modules estimate spatial transforms. Second, QCS evaluates each candidate alignment and decomposes the result into interpretable failure signals. Third, when the current alignment is insufficient, an LLM-based decision agent interprets the QCS decomposition and pipeline metadata to select an appropriate recovery strategy family rather than estimating image transformations directly. Fourth, an optional human-guided experience library stores user-validated recovery strategies for future reuse. This optional path is not required for routine automated registration; it provides an interface for users to convert validated refinements into reusable strategy rules. Operationally, the workflow proceeds as a four-stage cascade: input modality classification, routing to the corresponding deterministic pipeline, QCS-based evaluation and return of the best-scoring acceptable result. If the initial cascade does not reach the task-specific QCS threshold, the autonomous rescue loop is entered. The agent selects a named strategy from a predefined library, such as expanded orientation search, alternative preprocessing, channel re-selection, ROI-scale registration or non-rigid recipe escalation. The selected strategy is then executed by the deterministic backend and re-evaluated by QCS. Thus, the agentic layer does not replace exhaustive parameter search or low-level registration optimization. Instead, it performs failure-mode diagnosis and strategy routing, separating numerical transform estimation from high-level recovery selection and avoiding the treatment of heterogeneous recovery actions as an undifferentiated exhaustive search space.

### Shared registration core and transform model

All modality-specific pipelines share a common transform composition, moving from discrete orientation and scale hypotheses to affine, sub-pixel and optional non-rigid refinement. For image-based tasks, the transform can be written as a composition of a scale hypothesis *S_ρ_*, a flip *F*, a rotation *R_θ_*, an affine matrix *A* and an optional non-rigid deformation *ϕ*:

*T* =*ϕ∘ A ∘ R_θ_ ∘ F ∘ S_ρ_.*Here, *S_ρ_* represents the coarse pixel-ratio or scale correction, F is the flip hypothesis, *R_θ_* is the rotation hypothesis, A is the affine correction, and *ϕ* is the optional non-rigid deformation. The composition symbol ∘ means that transforms are applied from right to left: scale first, then flip, rotation, affine correction, and finally local deformation.

For two-dimensional affine registration, coordinates are represented in homogeneous form:

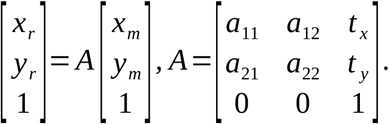

Here, (*x_m_*, *y_m_*) and (*x_r_*, *y_r_*) denote corresponding coordinates in the moving image and the H&E reference image, respectively, and A denotes a 3 × 3 homogeneous affine matrix. The submatrix *a_ij_* encodes rotation, scaling and shear, whereas *t _x_* and *t _y_* encode translation along the horizontal and vertical axes. In practical terms, this affine transform accounts for global differences in position, orientation, size and mild tissue shear before local deformation refinement.

Images were read using OpenSlide^43^ at a registration resolution of 8.0 micrometres per pixel, choosing the pyramid level whose microns-per-pixel value was closest to the target resolution. For plain TIFF files without embedded resolution metadata, the longest edge was downsampled to approximately 4000 pixels. Cross-modal preprocessing converted H&E to a hematoxylin-emphasizing grayscale representation using HED color deconvolution^44^ followed by 1st–99th percentile normalization to an 8-bit intensity range. This representation emphasizes the nuclear-density information shared by hematoxylin and DAPI while suppressing modality-specific color differences. Rigid and similarity transforms were optimized with Mattes mutual information^45^ using multi-resolution pyramids. Mutual information and normalized mutual information were defined from a joint intensity histogram of the fixed and warped moving images:

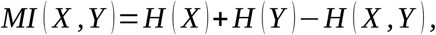

Here, *H* (*X*) and *H* (*Y*) are the entropy of the two single-image intensity distributions, *H* (*X, Y*) is their joint entropy, and *MI* (*X, Y*) measures how much information the two images share. Higher MI indicates stronger statistical dependence after registration.

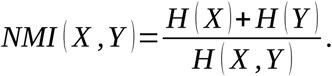

Here, NMI^46^ is a normalized version of mutual information that is less sensitive to the absolute intensity scale of the two modalities. It is useful for DAPI-to-H&E and IHC-to-H&E registration because the modalities do not share the same raw intensity appearance.

Similarity registration used multi-initialization to avoid poor local optima: moment-based initialization, geometry-based initialization and orthogonal rotation offsets were all tested, and the initialization with the best NMI was selected. For intensity-inverted modalities such as IHC, an additional shape-based similarity variant initialized from Otsu-thresholded^47^ tissue masks with seven rotation candidates. Affine refinement was then applied on the pre-aligned image using the same metric and optimizer. Optional non-rigid registration used either a BSpline^48^ transform or a dense displacement field. The resulting deformation map was written as:

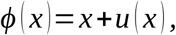

where *ϕ* (*x*) denotes the deformation map, *x* is an image coordinate, and *u* (*x*) is the local displacement vector at *x*. This formulation allows local tissue regions to bend or shift after global alignment.

For large whole-slide images, full-resolution output was generated by tile-streaming the moving image through the composed transform using 1024 *×* 1024-pixel tiles, bilinear interpolation and white border padding, and was written as a deflate-compressed OME-TIFF. For large non-rigid images, a tile-based displacement variant processed 2048 *×* 2048-pixel tiles with 256-pixel overlap and blended per-tile displacement fields using Hanning windows. A boundary-focused BSpline variant restricted optimization to a tissue-boundary ring when edge distortion needed correction without disturbing already-aligned tissue interiors.

### Composite Quality Score (QCS)

QCS is the central acceptance functional in ACCREDIT. It evaluates each registration attempt and determines whether the result should be accepted, rescued or rejected. The score is designed to reject candidates that appear plausible under one metric but fail under another, such as high tissue overlap with poor boundary agreement, high NMI in a homogeneous region despite gross offset, or warped tissue placed on H&E background. Given a warped moving mask and a reference H&E tissue mask, tissue Dice^49^ is defined as *M_m_^T^, M_r_*

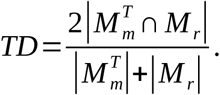

Here, TD is Tissue Dice, *M_m_^T^* is the warped moving tissue mask and *M_r_*is the H&E tissue mask. The numerator is the shared tissue area after registration, and the denominator normalizes by the total tissue area in both masks. TD is high when the overall tissue footprints overlap well.

Boundary Dice is computed on a narrow band around the tissue contour. Let *∂ M* be the tissue boundary and *B_r_* (*M*)=*dilate* (*∂ M, r*) be a boundary band of radius *r* pixels; *r* =8 pixels were used for QCS evaluation unless otherwise stated.

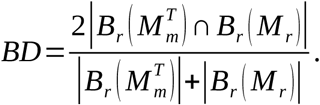

Here, BD is Boundary Dice, and *B_r_* (*M*) is a narrow band around the tissue boundary of mask M with radius *r*. BD focuses on contour agreement rather than bulk tissue overlap, so it is sensitive to small boundary-level shifts that TD may miss.

Hausdorff distance^50^ measures the largest boundary discrepancy between the warped moving tissue contour and the H&E reference contour. Let *C_m_^T^* and *C_r_* be contour point sets extracted from the warped moving and reference masks. The symmetric Hausdorff distance is

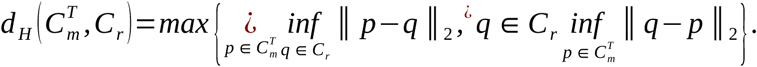

Here, *C_m_^T^* and *C_r_* are the boundary point sets of the warped moving tissue and reference H&E tissue. For each point on one boundary, the formula finds the nearest point on the other boundary; the Hausdorff distance is the largest of these nearest-neighbor distances in either direction. Thus, *d _H_* measures the worst boundary mismatch, not the average mismatch.

Because lower boundary distance indicates better alignment, the raw distance is converted into a bounded Hausdorff distance score using a resolution-dependent ceiling *H _max_*=500 *× L_eval_* / 4000, where *L_eval_* is the longest edge of the evaluation image in pixels:

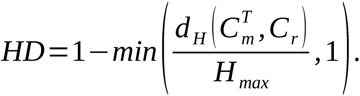

Here, *H_max_*is the resolution-dependent maximum tolerated boundary error. The raw distance *d _H_* is converted into a high-is-better HD score: HD is close to 1 when the largest boundary mismatch is small, and approaches 0 when the mismatch reaches or exceeds *H_max_*.

This distinction keeps the mathematical definition consistent with the implementation while allowing the QCS formula to use a high-is-better score.

Feature consistency was measured using the ORB^51^ inlier ratio after RANSAC^52^. If *N_match_* is the number of tentative feature matches and *N_inlier_*is the number retained by RANSAC, the inlier score is saturated at 0.15 to prevent this component from dominating QCS:

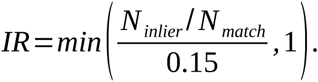

Here, *N _match_*is the number of tentative ORB feature matches and *N _inlier_* is the subset retained by RANSAC after registration. The ORB/RANSAC inlier ratio was saturated at 0.15 before normalization, converting values at or above 0.15 to the maximum feature-consistency score. This saturation prevents the feature-consistency term from disproportionately influencing the total QCS when the inlier ratio is high. NMI was computed from a 64-bin joint histogram and mapped to [0, 1] using a linear saturation around the empirical range observed in cross-modal registration:

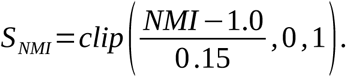

Here, *S_NMI_*is the bounded NMI score used inside QCS. The clip operation truncates values to [0, 1].

Two multiplicative penalties suppress degenerate solutions. The first penalizes under-coverage when the warped moving tissue covers less than 20% of the H&E tissue. The second penalizes warped moving signal that falls on H&E background:

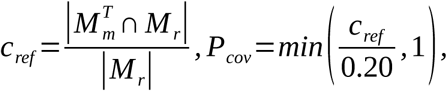

Here, *c_ref_*is the fraction of H&E tissue area covered by the warped moving tissue. *P_cov_* penalizes cases where the warped moving image covers too little of the H&E tissue, which is important for partial-field-of-view and degenerate-corner failures.

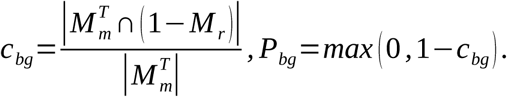

Here, *c_bg_*is the fraction of warped moving tissue that falls on H&E background. *P_bg_* reduces the score when the moving tissue is placed outside the reference tissue area, even if another metric appears superficially strong. The final QCS is a weighted sum of five bounded component scores multiplied by the two penalties:

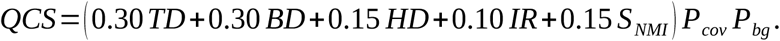

Here, TD, BD, HD, IR and *S_NMI_* are the five bounded quality components, and *P_cov_* and *P_bg_* are multiplicative penalties. The weights make tissue overlap and boundary agreement the dominant terms, while feature consistency and cross-modal intensity similarity provide supporting evidence.

All component scores were clamped to [0,1]. A QCS of at least 0.65 was classified as good and accepted without rescue. Scores from 0.45 to 0.65 were considered acceptable but eligible for rescue. Scores below 0.45 were considered poor and triggered rescue, and scores below 0.25 were classified as failed. A silent failure was operationally defined as a registration run that completed without error but produced QCS < 0.25, indicating gross misregistration. The thresholds provide a general interpretive scale for QCS, whereas individual pipelines may additionally apply task-specific escalation gates tuned to the residual structure of each registration task.

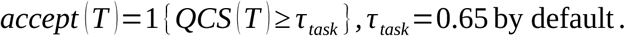

Here, *accept* (*T*) is a binary decision rule for whether a transform is accepted. *τ _task_* is the task-specific QCS threshold; 0.65 is the default threshold, but individual pipelines can apply stricter or additional gates when needed.

## Adaptive rescue and strategy-learning loop

### LLM-guided rescue agent

The autonomous rescue agent was invoked when the initial modality-specific cascade produced a QCS below the acceptance threshold or when the output was classified as borderline by a task-specific quality gate. The agent is formulated as a policy over a finite strategy library rather than as a registration optimizer. For each attempt t, the system records a diagnostic state containing the total QCS, individual component scores, coverage penalties, modality type, image-scale metadata, transform history and the configuration used in the previous attempt:

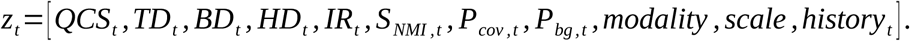

Here, *z_t_*denotes the diagnostic state observed after attempt t. The entries include the total quality score, the five bounded QCS components, the two multiplicative penalties, modality metadata, image-scale information, the transform history and the previous pipeline configuration. This representation makes the rescue step explicit: the agent reasons over measurable failure signals and pipeline metadata, rather than directly over unstructured raw images.

Based on this diagnostic record, the agent infers the likely failure mode, such as incorrect orientation, insufficient contrast normalization, partial field-of-view overlap, modality-descriptor mismatch or residual local deformation. It then selects an alternative strategy from a predefined modality-specific library. For DAPI-to-H&E registration, available strategies included expanded orientation search, alternative DAPI channel selection, MIP/CLAHE^53^ preprocessing, gamma enhancement and contour-chain backup registration. For CODEX-to-H&E registration, strategies included nuclear-channel rescue, protein-channel reselection and ROI-scale search. For IHC-to-H&E registration, strategies included alternative affine re-estimation and non-rigid recipe escalation. The strategy-selection step was written as a policy acting on the unused subset of the strategy library:

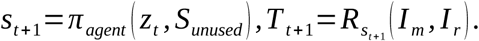

Here, *π _agent_*denotes the rescue-agent policy, *S_unused_* is the set of recovery strategies that have not yet been attempted, and *R_s_* denotes the deterministic registration module or parameter recipe associated with strategy s. The selected strategy *s_t_* is therefore a named and auditable registration action, while the numerical transform itself is still estimated by the deterministic backend.

Each rescue attempt was executed by the deterministic registration backend and re-evaluated by QCS. The agent did not directly modify image pixels or estimate transforms. Instead, it selected which validated registration module and parameter set should be attempted next.

The loop terminated when the task-specific acceptance threshold was reached, when no remaining strategy improved QCS by a minimum margin, or when the maximum number of rescue attempts was reached. The final output was selected as the highest-QCS registration among all attempted configurations, not necessarily the last attempted result:

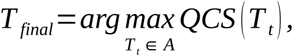

Here, *A*={*T* ₀ *, T* ₁ *,…, T _t_* }is the set of all transforms attempted during the initial cascade and subsequent rescue iterations.

### Optional user-guided strategy learning

ACCREDIT includes an optional user-guided strategy-learning path for cases in which the output from automated registration and autonomous rescue requires further refinement or user validation. This path is not required for routine automated registration. Instead, it provides an interface for user-initiated correction, validation and strategy learning. When a user inspects a difficult case, requests further refinement, provides a correction strategy or validates a successful rescue configuration, the system records the diagnostic state before refinement, the successful strategy sequence, the resulting quality improvement and the retrieval condition under which the strategy should be reused. Each stored experience record is represented as:

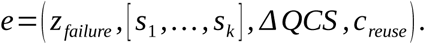

Here, e denotes a stored experience record. The term *z_failure_* denotes the diagnostic state of the failure case, [*s* ¿¿ 1,…*, s* ¿¿ *k*]¿¿ denotes the successful sequence of strategy steps, *ΔQCS* denotes the observed quality improvement, and *c_reuse_* denotes the condition under which the rule should be retrieved for future samples. Thus, the knowledge base stores not only the final transform but also the failure signature and the reasoning path that made the correction reusable.

At runtime, the experience knowledge base is queried for stored records whose diagnostic signatures are similar to the current failure state. Retrieved rules can be prioritized before generic rescue strategies, allowing previously validated refinements to be reused while keeping transformation estimation deterministic. Thus, optional user feedback is converted into reusable pipeline-level knowledge rather than being treated as a one-time manual correction.

### Modality-specific pipelines and downstream analysis

Detailed descriptions of the four modality-specific registration pipelines — the DAPI-to-H&E pipeline, the CODEX-to-H&E pipeline, the cell-boundary-to-H&E pipeline and the IHC-to-H&E hybrid backend — are provided in Supplementary Methods. Each pipeline implements the shared registration core and QCS-gated acceptance described above, with modality-specific channel selection, initialization and refinement strategies. Downstream cross-modal analysis procedures, including identification of cellular neighborhoods from CODEX protein data, projection of CODEX-defined cellular neighborhoods to Xenium space, TCGA-COAD bulk RNA-seq deconvolution, secondary follicle-like niche analysis and ACCREDIT-enabled single-cell H&E labeling, are also detailed in Supplementary Methods.

## Data Availability

### Xenium DAPI-to-H&E benchmark

We assembled 16 publicly available Xenium datasets from 10x Genomics, spanning 13 tissue types, including breast, cervical, lung, ovary, mouse brain, bone marrow, liver, colorectal, kidney, pancreas, skin, heart and prostate. The benchmark included both FFPE and fresh-frozen preparations, and included two instrument generations: Xenium V1 versions 1.0.1–2.0.0 and Xenium Prime version 3.0.0, with one protein-panel dataset at version 4.0.0. Each sample provided a matched H&E whole-slide image, a multi-channel DAPI morphology image, an official 10x alignment file and nucleus/cell boundary coordinates in Parquet format. Dataset identifiers and download information are listed in Supplementary Table.

### Glioma CODEX DAPI-to-H&E benchmark

To assess cross-platform generalization beyond Xenium morphology images, we included a glioma spatial-omics benchmark consisting of 12 matched CODEX DAPI and H&E image pairs. In this dataset, the CODEX DAPI channel captures nuclear morphology from a multiplexed protein-imaging workflow, whereas the matched H&E section provides conventional histological morphology. The CODEX DAPI image was registered to the corresponding H&E image using the DAPI-to-H&E pipeline, and registration quality was evaluated using the same QCS protocol as the Xenium benchmark. This dataset was analyzed together with the 16 Xenium DAPI-to-H&E samples to form a 28-sample combined DAPI-to-H&E benchmark used for the distribution-level analysis in **Supplementary Fig. 2**.

### Representative PASC IHC-to-H&E samples

Two pairs of adjacent pancreatic adenosquamous carcinoma serial sections obtained from surgically resected specimens were stained with mouse monoclonal p40 antibody (BC28, BioCare Medical, Pacheco, CA) IHC and H&E. Each pair was digitized as a whole-slide image at 40x magnification. These samples were used as representative IHC-to-H&E registration examples in **Fig. 4**. The samples are not publicly available because of institutional data-sharing restrictions. Access may be granted upon reasonable request to the corresponding author. The representative pancreatic adenosquamous carcinoma IHC-to-H&E samples were retrospectively obtained from de-identified archival specimens under a protocol with a Health Insurance Portability and Accountability Act (HIPAA) waiver approved by the institutional review board at Oregon Health & Science University (IRB#23756).

### CODEX/Xenium dataset for downstream cross-modal analysis

The downstream cross-modal analysis in **Fig. 5** used adjacent Xenium and CODEX sections linked through the same H&E-anchored coordinate system. The Xenium section provided transcriptomic profiles, and the CODEX section provided protein measurements and protein-defined cellular neighborhoods. Physical alignment between sections established the shared spatial frame used for cellular-neighborhood transfer, comparative cell-type composition analysis and secondary follicle-like niche interpretation.

## Code Availability

The source code for ACCREDIT is publicly available on GitHub at https://github.com/LeeZhou-bearway/ACCREDIT.

## Supporting information

Supplementary Information for ACCREDIT

## Acknowledgements

This work was supported by the following funding sources: the Department of Defense Prostate Cancer Data Science Award HT94252410551, NIH grants R01GM147365 and R01CA283171, and a CPRIT Scholar Award RR260021 from the Cancer Prevention and Research Institute of Texas (to Z. Xia); the Department of Defense Pancreatic Cancer Award HT94252510959, Kuni Foundation, and Knight Pilot Award CBTOP-2023-002 (to A. Kardosh); and NIH grants U01CA294548, U01CA278923 and R01CA186241 (to R.C. Sears). The research reported here used computational infrastructure supported by the Office of Research Infrastructure Programs, Office of the Director, of the National Institutes of Health under Award Number S10OD034224. We thank the Brenden-Colson Center for Pancreatic Care (Dove Keith) and Knight BioLibrary (Aletha Lesch) for assistance in obtaining pancreatic tumor samples, as well as OHSU Histopathology Shared Resource (RRID: SCR_009977) for assistance with tissue sectioning, H&E and IHC staining (Kate Rice, Joscelyn Zarceno). The content is solely the responsibility of the authors and does not necessarily represent the official views of the funding agencies.

## Author Contributions

Z.X., L.Z. and T.R. conceived the idea. L.Z. led the study and developed the overall algorithmic architecture. T.R. developed the contour-registration algorithm incorporated into the cell-boundary-to-H&E pipeline. F.Z. performed the biological analyses. A.K., S.M.G., B.L., T.Z., C.T., Y.C., R.C.S and G.B.M. contributed to biological interpretation, data coordination, and evaluation of registration outputs. Z.X. supervised the study. L.Z., F.Z., T.R. and Z.X. wrote the manuscript with input from all authors. All authors reviewed and approved the final manuscript.

