## Supplementary Information for ACCREDIT for "ACCREDIT: A Quality-Aware Agentic Engine for Cell-resolved Cross-modal Image Registration with Dynamic Iterative Tuning"

#### DAPI-to-H&E pipeline

The DAPI-to-H&E pipeline registers fluorescence morphology images to matched H&E sections through a five-step QCS-gated cascade. The first step explicitly searches orientation and pixel-ratio hypotheses because DAPI and H&E inputs may differ by scanner-induced flips, 90-degree rotations and scale changes. The discrete candidate set is defined as:

$$\mathcal{C} = \mathcal{F} \times \mathcal{R} \times \mathcal{P},$$

$$|\mathcal{F}| = 4,\ |\mathcal{R}| = 4,\ |\mathcal{P}| = 6,\ |\mathcal{C}| = 96.$$

Here $\mathcal{F} = \{\text{none}, \text{vertical}, \text{horizontal}, \text{both}\}$ is the flip set, $\mathcal{R} = \{0°, 90°, 180°, 270°\}$ is the rotation set and $\mathcal{P} = \{0.5,0.7,0.8,1.0,1.2,1.5\}$ is the pixel-ratio set. Each candidate $c$ defines a coarse transform $T_c$. Candidate selection is written as a QCS-maximization problem at 2,000-pixel scan resolution:

$$c^* = \underset{c \in \mathcal{C}}{\operatorname{argmax}} \text{QCS}(T_c).$$

Here, $c^*$ denotes the selected coarse orientation-scale candidate, the flip, rotation and pixel-ratio combination whose coarse transform achieves the highest QCS among all candidates in $C$. For full-coverage samples with known pixel ratio, the search was reduced to 16 flip-rotation candidates. If the best candidate after the first scan had QCS < 0.60, the scan was repeated with maximum-intensity projection plus CLAHE preprocessing and with single-channel CLAHE preprocessing. The globally best candidate across all preprocessing variants was retained. This step is the main mechanism by which ACCREDIT avoids silent mirror or rotation failures before fine registration begins.

Step 2 refined the winning orientation with SimpleITK Similarity2D[1] registration using Mattes mutual information, 64 histogram bins, $30\%$ random sampling, Regular Step Gradient Descent, a learning rate of 1.0, a minimum step of 0.001, 1000 iterations and a $[4, 2, 1]$image pyramid at 0.25x resolution. Both moment-based and geometry-based initializations were tested to reduce dependence on a single starting point.

Step 3 performed NCC-guided angle, translation and optional anisotropic scale fine-tuning. Around the current angle, the pipeline swept $\pm 50$ degrees in 1-degree increments and then performed a translation grid search over $\pm 15$ pixels with 3-pixel step size. For samples requiring anisotropic scale correction, independent x and y scale factors were searched within $\pm 20\%$ of the dimension-based ratio at 1% increments, using gamma-corrected fluorescence images. NCC[2] was defined as:

$$\text{NCC}(X,Y) = \frac{\sum_x (X(x) - \bar{X})\,(Y(x) - \bar{Y})}{\sqrt{\sum_x (X(x) - \bar{X})^2 \sum_x (Y(x) - \bar{Y})^2}}.$$

Here, NCC is normalized cross-correlation, X and Y are local image patches, and bars denote mean intensity within the patch. NCC measures whether two patches have similar spatial intensity patterns after subtracting their means.

Step 4 applied OpenCV ECC[3] refinement using a Euclidean motion model with 500 iterations, convergence tolerance $1 * 10^{-6}$and Gaussian filter size 5 at $0.5\text{x}$ resolution. Step 5 applied BSpline elastic deformation with an $8\text{x}8$ control-point mesh, Mattes mutual information, 50 bins, $20\%$ regular sampling, L-BFGS-B[4] optimization, 200 iterations and a $[2, 1]$pyramid. Displacement magnitudes exceeding the median plus two standard deviations within the tissue region were clamped to prevent boundary over-stretching:

$$u_{\text{clamped}}(x) = u(x) \cdot \min\left(1, \frac{\text{median}(|u|) + 2\sigma(|u|)}{|u(x)|}\right).$$

Here, u(x) is the estimated local displacement vector, and ||u(x)|| is its magnitude. The median and standard deviation are computed over displacement magnitudes inside the tissue region. Displacements larger than this robust threshold are clamped, preventing unrealistic local stretching at tissue boundaries.

If final QCS remained below 0.65, the agent could re-invoke the pipeline with alternative channel selection, gamma enhancement or a contour-chain backup strategy. Each re-run was evaluated by QCS and accepted only if it improved the best score or exceeded the task acceptance threshold.

**CODEX-to-H&E pipeline**

The CODEX-to-H&E pipeline registers multi-channel protein-fluorescence images to H&E through a four-step QCS-gated cascade. Registration of IHC-stained adjacent serial sections presents a distinct deformation challenge and is therefore described separately in "IHC-to-H&E hybrid backend". Because the most informative CODEX channel varies by sample, channel selection was formalized as either a channel-coverage-guided screening step or a QCS-guided selection step, depending on whether candidate registrations were already available.

For each available CODEX channel $k$ , a tissue-like foreground mask $M_k$ was extracted after intensity normalization and foreground thresholding. We defined a channel coverage score,

$$C_{ch(k)} = \frac{|M_k|}{|M_r|},$$

where $M_k$ is the tissue-like foreground mask extracted from channel $k$, and $M_r$ is the H&E reference tissue mask. This score measures the relative amount of tissue-like signal available in each CODEX channel and was used only for channel screening. It is distinct from the QCS coverage penalties defined in "Composite Quality Score", which evaluate the spatial relationship between a warped moving mask and the H&E reference mask after registration. Initial channel screening selected the channel with the highest channel coverage score:

$$k^* = \arg\max_k C_{ch(k)}, k = 1, \dots, K.$$

Here $K$ is the set of available CODEX channels. When candidate registrations were available, channel selection was instead based on the full QCS:

$$k^* = \arg\max_k QCS(T_k)\,, k = 1, \ldots, K.$$

where $T_k$ is the candidate transform estimated using channel $k$. Thus, channel-coverage-guided selection was used before reliable registration candidates were available, whereas QCS-guided selection was used after candidate transforms had been estimated.

Step 1 registered the best-coverage protein channel to H&E using the same NMI similarity registration, NCC angle fine-tuning, ECC refinement and BSpline deformation modules as the DAPI-to-H&E pipeline. Step 2 always ran a parallel nuclear-channel rescue on channel 0 (DAPI), using multi-initialization NMI, NCC rotation scanning on downsampled tissue masks, phase correlation[5] for sub-pixel translation and ECC polishing. The Step 1 and Step 2 results were compared by QCS, and the better result was retained:

$$T_{keep} \in \{T_{protein}, T_{nuclear}\},$$

$$QCS(T_{keep}) = \max\{QCS(T_{protein}), QCS(T_{nuclear})\}$$

Here, $T_{\text{protein}}$ and $T_{\text{nuclear}}$ are the candidate transforms produced from protein-channel and DAPI/nuclear-channel registration, respectively. The pipeline keeps the transform with the higher QCS, allowing nuclear morphology to rescue weak protein-channel alignment.

If QCS remained below 0.60, Step 3 re-registered the protein channel with a wider rotation scan and phase-correlation translation recovery. If QCS remained below 0.35, indicating a large field-of-view or scale mismatch, Step 4 performed ROI-scale registration over pixel-ratio hypotheses [1.0,1.5,2.0,2.5,3.0,3.5,4.0] with per-ratio NMI ranking. Decision thresholds were QCS ≥ 0.60 for completion, QCS < 0.60 for protein-channel rescue, QCS < 0.35 for ROI-scale registration and QCS < 0.25 for user-guidance suggestions rather than silent acceptance.

**Cell-boundary-to-H&E pipeline**

The cell-boundary pipeline aligns Xenium per-cell polygon coordinates to the matched H&E section through a four-stage workflow and maps contour coordinates directly into H&E image space. This direction is important because it allows aligned polygons to be inspected against tissue morphology and used for downstream image-based analyses. On the 16-sample benchmark, the pipeline ran in a mean of 4.8 min per sample (range 1–10 min). Let a polygon vertex be represented in micrometres and let r be the H&E microns-per-pixel value. The initial pixel-coordinate conversion is:

$$v_i^{\text{px}} = \left(\frac{x_i^{\mu m}}{r}, \frac{y_i^{\mu m}}{r}\right).$$

Here, $v_i^{\mu m}$ is a polygon vertex in physical micrometre coordinates, r is the H&E microns-per-pixel value, and $v_i^{px}$ is the corresponding H&E pixel coordinate. This conversion places Xenium cell-boundary coordinates into the image coordinate system used by the H&E slide.

Stage 1 rasterized nucleus or cell polygons into a filled binary contour mask. In parallel, the H&E reference image was converted into a binary morphology image using a weighted RGB combination, contrast enhancement and Otsu[6] thresholding. Stage 2 estimated a global affine transform by aligning the contour mask to the H&E binary image using ORB feature matching at reduced scale, followed by affine estimation with RANSAC. If I is the RANSAC inlier set, the affine was estimated by minimizing inlier reprojection error:

$$A^* = \underset{A}{\operatorname{argmin}} \sum_{(p_i,q_i)\in\mathcal{I}} \parallel q_i - Ap_i \parallel_2^2.$$

Here, $p_i$ and $q_i$ are matched points from the contour-derived moving mask and the H&E morphology mask. A* is the affine transform that minimizes squared correspondence error over the RANSAC inlier set I.

Stage 3 refined the globally aligned contour mask on a $32 \times 32$ block grid. For each sufficiently populated block, template matching estimated a local displacement relative to the H&E reference. Sparse or low-confidence blocks were rejected or merged, and local updates were accepted only when they improved block-level overlap. Accepted displacements were smoothed into a continuous local field:

$$u_b^* = \underset{u\in\mathcal{U}_b}{\operatorname{argmax}} \mathrm{NCC}\big(C_b(x+u), H_b(x)\big),$$

Here, $b$ indexes a local block, $C_b$ is the contour-density image in that block, $H_b$ is the corresponding H&E morphology patch, and $U_b$ is the set of candidate local shifts. The selected displacement $u_b^*$ is the shift with the highest NCC score. The accepted sparse block displacements were placed on the block grid. Missing or rejected blocks were filled by interpolation from neighboring accepted blocks, and the resulting displacement map was smoothed to form a continuous displacement field $u(x)$[7]. The final boundary-to-H&E mapping was then defined as:

$$T_{\mathrm{boundary}}(x) = A^*x + u(A^*x).$$

Here, $T_{boundary}$ denotes the final transformation from cell-boundary coordinates to H&E image coordinates, $A^*$ is the selected global affine transform, and $u(A^*x)$ is the local displacement evaluated after affine alignment. This maps each boundary coordinate into H&E image space while correcting residual local offsets.

Stage 4 applied the composed global affine and local displacement field to every polygon vertex and exported the transformed polygons as GeoJSON. Final quality was quantified by global Dice and interior Dice between the warped contour mask and the H&E binary morphology mask. When global Dice remained below 0.50, an optional rescue module switched from sparse feature matching to dense signal correlation by comparing an H&E-derived nuclear channel with a blurred contour-density map through NCC rotation scanning and phase-correlation translation recovery. If Dice remained below 0.20 after rescue, the pipeline reported the failure state and provided optional guidance prompts rather than silently accepting the result.

**IHC-to-H&E hybrid backend**

Registering adjacent serial sections stained with IHC and H&E requires handling both cross-modal appearance mismatch and non-rigid tissue deformation introduced during sectioning, mounting and staining. We therefore implemented an ACCREDIT-IHC hybrid backend that combines feature-based global initialization, iterative non-rigid deformation and final Demons polishing within the same QCS-gated framework used by the other ACCREDIT pipelines.

First, pretrained SuperPoint[8] and LightGlue features[9], used with upstream weights and without histology-specific fine-tuning, were matched between the IHC-derived intensity image and the H&E reference image. A global affine transform was then estimated from matched keypoints using RANSAC. Second, local tissue deformation was estimated using a DHR-inspired non-rigid registration module based on local normalized cross-correlation and diffusion regularization. Third, a Symmetric Forces Demons polish[10] implemented in SimpleITK was applied for 300 iterations to refine residual local offsets. The non-rigid deformation field was estimated by minimizing an objective of the form:

$$E(u) = -\mathrm{LNCC}(I_r, I_m \circ \varphi_u) + \lambda \mathrm{R}(u).$$

Here, $I_r$ is the H&E reference image, $I_m$ is the IHC-derived moving image, $u_{(x)}$ is the displacement field using the non-rigid deformation notation defined in "Shared registration core and transform model", LNCC denotes local normalized cross-correlation and $\mathrm{R}_{(u)}$ denotes diffusion regularization weighted by $\lambda$. The first term encourages local image similarity after warping, whereas the second term penalizes spatially abrupt deformation and reduces unrealistic folding or noisy displacement fields.

The fixed ACCREDIT-IHC hybrid recipe alternated three rounds of feature-based affine re-estimation and non-rigid deformation, followed by the 300-iteration Demons polish. This alternating design distinguishes the backend from a single affine initialization followed by one deformation step: repeated affine re-estimation corrects residual global drift that might otherwise be absorbed incorrectly by the deformation field. The overall transform was written as

$$C_i = \phi_i \circ A_i, \qquad i = 1,2,3\,,$$

$$T_{IHC} = D_{300} \circ C_3 \circ C_2 \circ C_1,$$

where $C_i = \phi_i \circ A_i$ denotes the affine and non-rigid refinement block at round $i$, $A_i$ is the affine transform, $\phi_i$ is the non-rigid deformation, $D_{300}$ is the final 300-iteration Demons polishing transform, and $\circ$ denotes transform composition applied from right to left.

The ACCREDIT-IHC hybrid backend returned immediately if QCS evaluated at 512-pixel resolution was at least 0.70. Otherwise, optional escalation stages tested alternative non-rigid configurations from the recipe registry or invoked the LLM rescue agent. Thus, this backend should not be interpreted as a new deformation model alone. Rather, it is a fixed ACCREDIT-IHC recipe that integrates DHR-inspired non-rigid registration, repeated affine correction, Demons polishing and QCS-gated escalation without per-sample tuning on the reported benchmark metric.

#### Downstream cross-modal analysis

##### Identification of cellular neighborhoods from CODEX protein data

Aligned CODEX cells with valid cell-type annotations were used for CN analysis. For each CODEX cell, we quantified the composition of its local microenvironment by identifying neighboring CODEX cells within an 80-μm radius[11] in the shared H&E coordinate system using a k-d tree-based nearest-neighbor search. Neighbor counts were summarized by annotated cell type and normalized to proportions, generating a cell-by-cell neighborhood composition matrix. This matrix was z-score standardized and subjected to principal component analysis (PCA), and the top principal components were used for unsupervised clustering with k-means[12]. Candidate values of k were evaluated over a range of 2-20 using silhouette score[13], Calinski-Harabasz[14] index, Davies-Bouldin[15] index, cluster-size stability, and biological interpretability, including the ability to resolve the TLS-like region. Based on these combined criteria, we selected k = 8 and defined eight CN states (CN1-CN8). Each CODEX cell was assigned a CN label according to its k-means cluster membership.

##### Projection of CODEX-defined CNs to Xenium space

To transfer CN labels from protein space to RNA space, we leveraged the shared post-alignment H&E coordinate system. After physical registration, CODEX and Xenium cells were represented in the same spatial reference frame. For each Xenium cell, we identified its nearest CODEX cell by Euclidean distance using a k-d tree and assigned the corresponding CODEX-derived CN label to that Xenium cell. The nearest-neighbor transfer distance was retained as a quality-control metric. This procedure enabled direct comparison of CN spatial distributions and cell-type compositions between CODEX protein data and Xenium RNA data. This label-transfer strategy allowed protein-defined spatial states to be interrogated in matched transcriptomic space without re-clustering Xenium cells independently. To characterize transcriptional programs associated with each CN, we applied COSG[16] to the Xenium dataset using mapped CN labels as the grouping variable. The top-ranked genes for each CN were used to define CN-associated marker sets.

##### TCGA-COAD bulk RNA-seq data processing and CN-based deconvolution

TCGA-COAD bulk RNA-seq data[17] were obtained from TCGAbiolinks R package (v.1.32.0). For CN-based deconvolution, Xenium cells with transferred CN annotations were used as the single-cell reference. MuSiC R package (v.1.0.0)[18] was applied to estimate CN proportions in bulk TCGA-COAD samples using shared genes between Xenium and bulk RNA-seq datasets. This analysis generated sample-level CN abundance profiles for downstream clinical association analyses. CN deconvolution scores were then integrated with TCGA-COAD clinical and survival data for prognostic evaluation. Overall survival (OS) was analyzed using univariate Cox[19] proportional hazards models.

#### Secondary follicle-like niche analysis

A secondary follicle-like (SFL-like) region was identified by visual inspection of the aligned H&E image based on lymphoid-like morphology. The region was partitioned into core and boundary compartments, and composition was analyzed separately in CODEX protein space and Xenium RNA space using the transferred CN labels and modality-specific cell-type annotations. This analysis tested whether the aligned modalities supported a coherent core-boundary organization within the immune niche.

#### ACCREDIT-enabled single-cell H&E labeling for downstream machine learning

To evaluate whether ACCREDIT-enabled cell-boundary registration could support downstream morphology-based machine learning, H&E image morphology was linked with Xenium-derived single-cell molecular labels. ACCREDIT registered cell-boundary and centroid information into the H&E coordinate system, defining single-cell-resolution H&E regions corresponding to individual Xenium cells. Cell-type labels derived from Xenium gene-expression profiles were assigned to epithelial, stromal and T/B-cell categories and transferred to the matched H&E single-cell regions. Each labeled H&E patch was encoded with a pretrained image encoder to obtain morphology-based features, which were then analyzed by k-means clustering with $k = 3$. Cluster assignments were compared with transferred Xenium labels using class-specific precision, recall and F1-score for epithelial, stromal and T/B-cell classes. This downstream application tests whether registration-derived single-cell H&E patches preserve cell-type-associated morphology structure and illustrates how accurate boundary registration can support morphology-based machine-learning workflows.

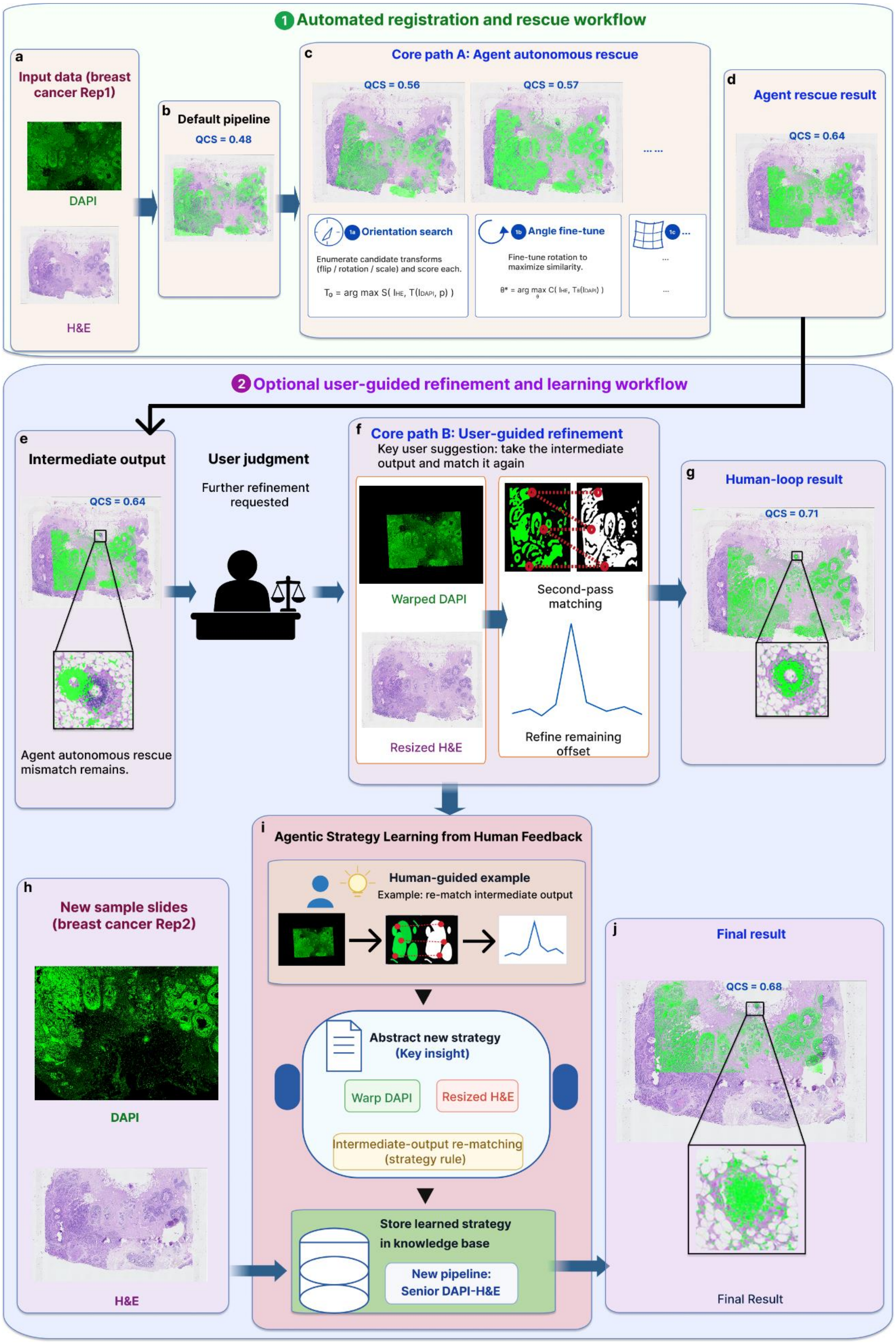

1 Automated registration and rescue workflow
a
Input data (breast cancer Rep1)
DAPI
H&E
b
Default pipeline
QCS = 0.48
c
Core path A: Agent autonomous rescue
QCS = 0.56
QCS = 0.57
... ...
1a Orientation search
Enumerate candidate transforms (flip / rotation / scale) and score each.
T0 = arg max S( IHE, T(IDAPI, p) )
1b Angle fine-tune
Fine-tune rotation to maximize similarity.
θ* = arg max C( IHE, Tθ(IDAPI) )
1c ...
...
...
d
Agent rescue result
QCS = 0.64
2 Optional user-guided refinement and learning workflow
e
Intermediate output
QCS = 0.64
Agent autonomous rescue mismatch remains.
User judgment
Further refinement requested
f
Core path B: User-guided refinement
Key user suggestion: take the intermediate output and match it again
Warped DAPI
Resized H&E
Second-pass matching
Refine remaining offset
g
Human-loop result
QCS = 0.71
h
New sample slides (breast cancer Rep2)
DAPI
H&E
i
Agentic Strategy Learning from Human Feedback
Human-guided example
Example: re-match intermediate output
Abstract new strategy
(Key insight)
Warp DAPI
Resized H&E
Intermediate-output re-matching (strategy rule)
Store learned strategy in knowledge base
New pipeline: Senior DAPI-H&E
j
Final result
QCS = 0.68
Final Result

***Supplementary Fig. 1* | *Automated registration and optional user-guided strategy learning for DAPI–H&E refinement.* *a–d,*** *Upper workflow showing the automated DAPI-to-H&E registration path using Breast Cancer FFPE Rep1 as an example. The input DAPI and H&E images are first processed by the default DAPI-to-H&E pipeline. When the initial output remains below the quality threshold, ACCREDIT enters the autonomous agentic rescue path, where orientation search, angle fine-tuning and non-rigid refinement are evaluated by QCS without user intervention. This automated workflow improves the alignment but may still leave residual local mismatch in difficult cases.* ***e–g,*** *Lower workflow showing an optional user-guided refinement interface for cases in which the user requests further improvement of the automated result. After inspecting the intermediate output, the user can provide guidance to reuse the warped DAPI image and resized H&E reference for second-pass matching. This optional refinement step corrects the remaining offset and improves local correspondence.* ***h–j,*** *The successful user-guided correction is abstracted into a reusable strategy rule, stored in the agent knowledge base and incorporated into an updated Senior DAPI-to-H&E pipeline. The learned strategy is then applied to a new Breast Cancer FFPE Rep2 case, demonstrating how optional user feedback can be converted into reusable pipeline-level knowledge for future automated registration.*

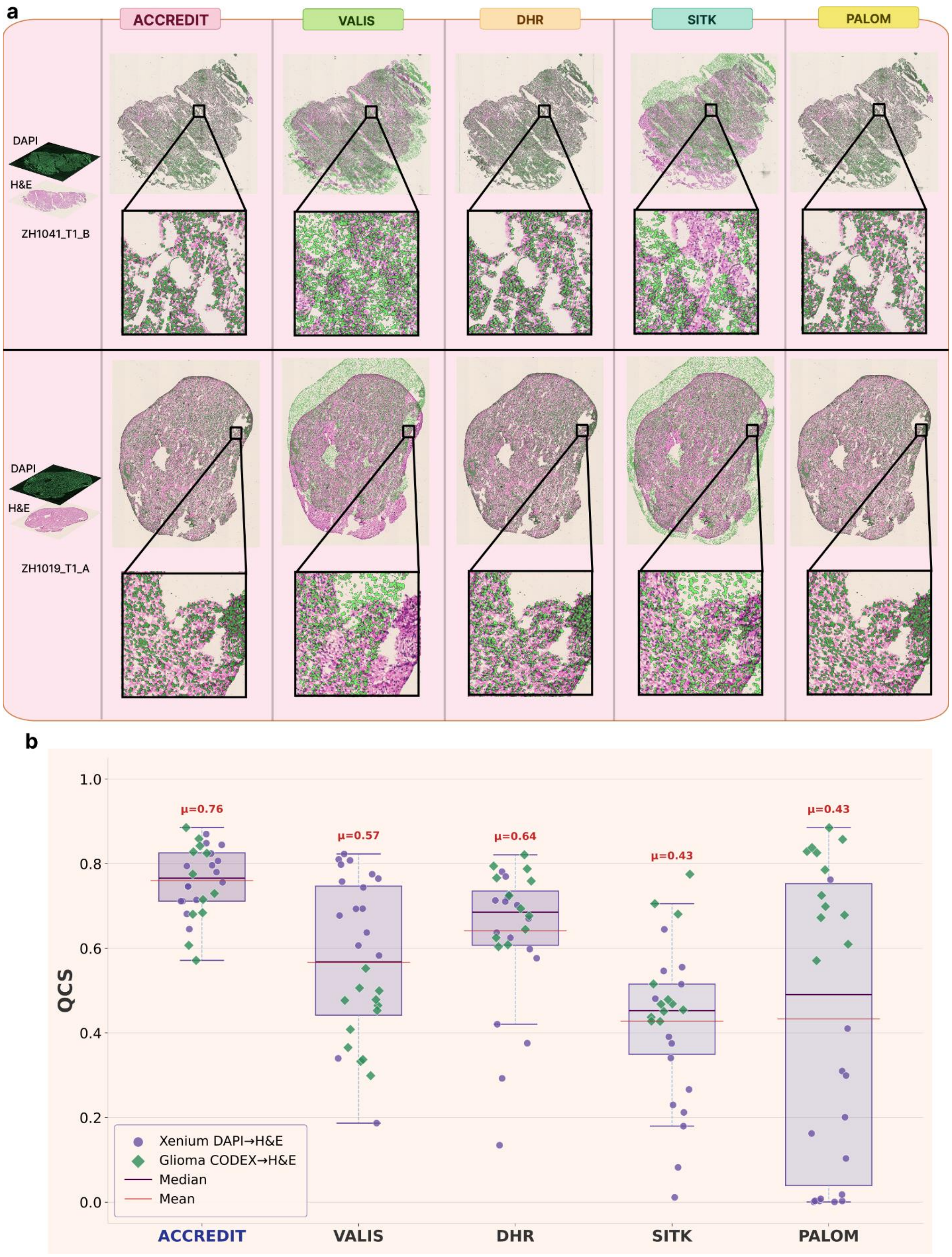

a
ACCREDIT
VALIS
DHR
SITK
PALOM
DAPI
H&E
ZH1041_T1_B
DAPI
H&E
ZH1019_T1_A
b
QCS
1.0
0.8
0.6
0.4
0.2
0.0
μ=0.76
μ=0.57
μ=0.64
μ=0.43
μ=0.43
Xenium DAPI→H&E
Glioma CODEX→H&E
Median
Mean
ACCREDIT
VALIS
DHR
SITK
PALOM

***Supplementary Fig. 2 | Combined DAPI–H&E registration benchmark across Xenium and glioma CODEX datasets. a**, Representative qualitative overlay comparison for glioma CODEX–H&E registration. Each row shows one glioma CODEX–H&E sample, with the input DAPI and matched H&E images shown on the left, followed by registration outputs from ACCREDIT and competing methods. DAPI signal is rendered in green and overlaid on the H&E image to visualize spatial correspondence. Zoomed regions highlight local alignment accuracy, residual mismatch, and deformation stability across methods. **b**, Distribution of composite quality scores (QCS) across five registration methods evaluated on 28 samples, including 16 Xenium DAPI→H&E pairs and 12 glioma CODEX→H&E pairs. Each point represents one sample; circles denote Xenium DAPI→H&E samples and diamonds denote glioma CODEX→H&E samples. Box plots show the median, interquartile range, and whiskers, with the mean QCS annotated above each method. ACCREDIT achieved the highest mean QCS ( $\mu$ = 0.76) and maintained a compact high-performance distribution across both datasets. In contrast, competing methods showed broader QCS distributions and more dataset-dependent variability, indicating less consistent registration performance across tissue types and imaging platforms.*

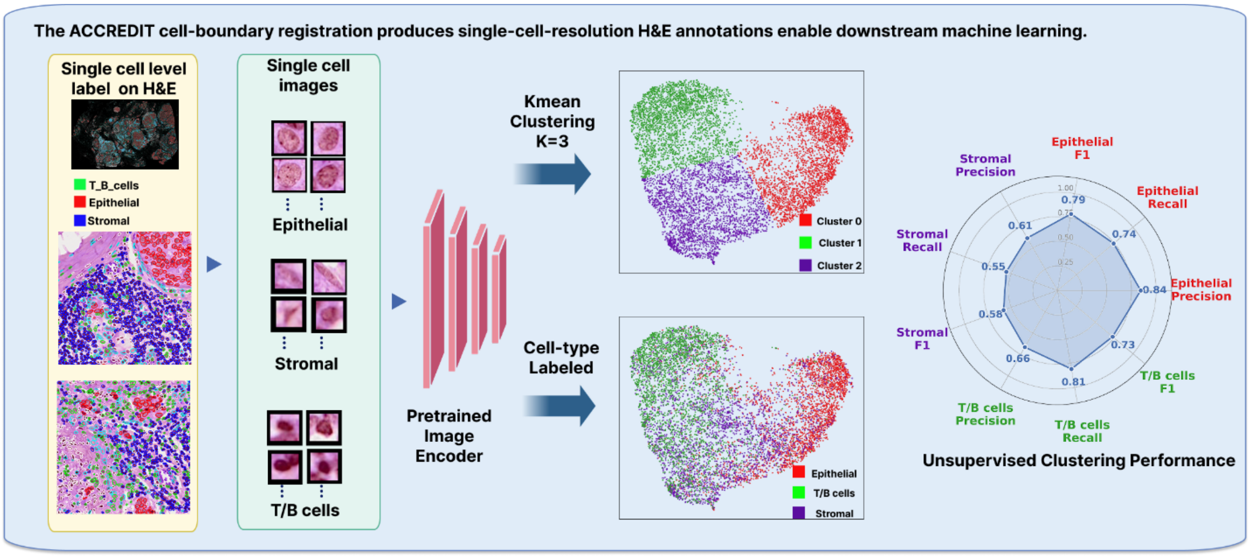


***Supplementary Fig. 3 | Single-cell H&E labeling enabled by ACCREDIT cell-boundary registration.***
*ACCREDIT registers cell-boundary information onto the corresponding H&E image, enabling single-cell-resolution H&E regions to be linked with Xenium-derived molecular profiles. Xenium provides gene-expression measurements for individual cells, from which cell-type labels are assigned. After registration, these labels can be transferred to the corresponding H&E single-cell patches, generating labeled morphology examples*

*for downstream machine-learning analysis. In this proof-of-concept workflow, H&E single-cell image patches were extracted for epithelial, stromal, and T/B-cell classes, encoded using a pretrained image encoder, and clustered with k-means at k = 3. The resulting morphology-based clusters were compared with Xenium-derived cell-type labels and evaluated using class-specific precision, recall, and F1-score.*

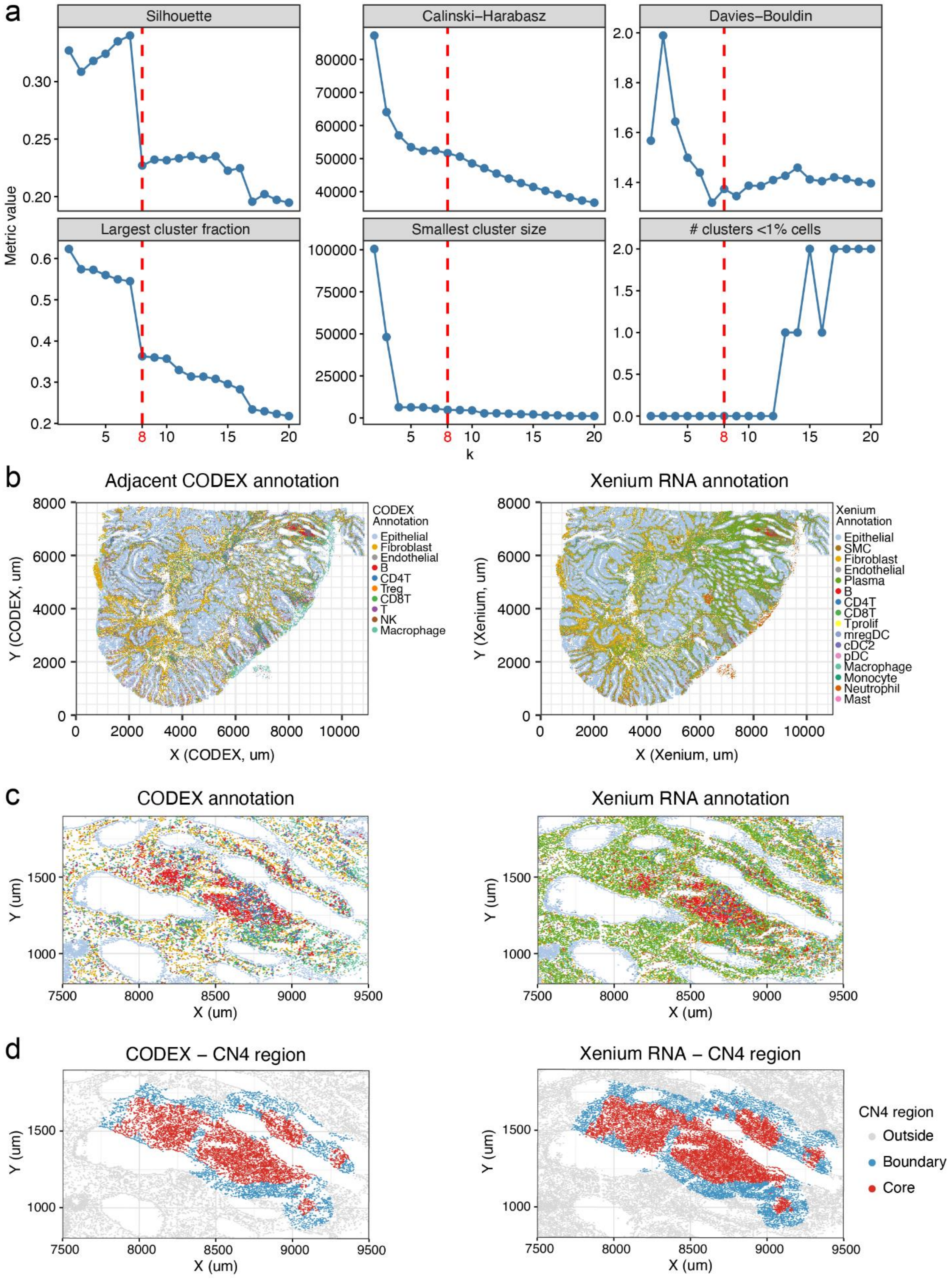
a
Silhouette
Calinski−Harabasz
Davies−Bouldin
Largest cluster fraction
Smallest cluster size
### clusters <1% cells
Metric value
k
b
Adjacent CODEX annotation
CODEX Annotation
Epithelial
Fibroblast
Endothelial
B
CD4T
Treg
CD8T
T
NK
Macrophage
Y (CODEX, um)
X (CODEX, um)
Xenium RNA annotation
Xenium Annotation
Epithelial
SMC
Fibroblast
Endothelial
Plasma
B
CD4T
CD8T
Tprolif
mregDC
cDC2
pDC
Macrophage
Monocyte
Neutrophil
Mast
Y (Xenium, um)
X (Xenium, um)
c
CODEX annotation
Xenium RNA annotation
Y (um)
X (um)
d
CODEX − CN4 region
Xenium RNA − CN4 region
CN4 region
Outside
Boundary
Core
Y (um)
X (um)

***Supplementary Fig. 4 Determination of CN number and validation of cross-modal cell-type annotation. a**, Evaluation of clustering performance across different values of k using multiple metrics, including Silhouette score, Calinski–Harabasz index, Davies–Bouldin index, largest cluster fraction, smallest cluster size, and number of clusters containing <1% of cells. k = 8 (red dashed line) was selected as an optimal balance between cluster stability and biological interpretability. **b**, Whole-section cell-type annotation of CODEX protein (left) and Xenium RNA data (right), demonstrating broadly consistent spatial distributions of major cell populations across modalities. **c**, Zoom-in views showing local concordance of cell-type annotation between CODEX and Xenium in representative regions. **d,** Spatial delineation of CN4 regions into core and boundary compartments in CODEX (left) and Xenium (right), illustrating consistent identification of structured immune niches across modalities.*

| Dataset Category | Source | Link / DOI | # Samples |
|---|---|---|---|
| CODEX Glioma | Zenodo | https://zenodo.org/records/12624860<br>DOI: 10.5281/zenodo.12624860 | 12 CODEX + 12 HE |
| Xenium DAPI | 10x Genomics | https://www.10xgenomics.com/datasets<br>CDN: cf.10xgenomics.com/samples/xenium/ | 14 datasets (HE + morphology + alignment) |
| DAPI-CODEX-HE | SPATCH Portal | http://spatch.pku-genomics.org/<br>DOI: 10.1038/s41467-025-64292-3<br>GitHub: github.com/zenglab-pku/SPATCH | 3 platforms x 3 tumors (Xenium5K, CosMx, VisiumHD) |

***Supplementary Table 1. Dataset sources and download links for the benchmark datasets used in ACCREDIT.***
